# Mechanism of outer kinetochore assembly on microtubules and its regulation by mitotic error correction

**DOI:** 10.1101/2023.07.20.549907

**Authors:** Kyle W. Muir, Christopher Batters, Tom Dendooven, Jing Yang, Ziguo Zhang, Alister Burt, David Barford

## Abstract

Kinetochores couple chromosomes to the mitotic spindle and transduce the energy of microtubule depolymerization to segregate the genome during cell division. Kinetochore – microtubule attachments are often initially erroneous and subject to a mitotic error correction (EC) mechanism that drives their turnover until biorientation is achieved. How this is accomplished and regulated, and how kinetochore-mediated chromosome segregation occurs at a molecular level remain major outstanding questions. Here we describe the cryo-electron microscopy (cryo-EM) structure of the budding yeast outer kinetochore Ndc80 and Dam1 ring complexes assembled onto microtubules. We observe coordinated interactions of the outer kinetochore complexes through multiple interfaces, in addition to a short staple within the Dam1 subunit that facilitates Dam1c ring assembly. Perturbation of these interfaces results in loss of yeast viability. Force-rupture assays indicated this is a consequence of substantial reductions in kinetochore – microtubule binding strength. EC-mediated phosphorylation of Ndc80c-Dam1c interfaces would drive complex disassembly, whereas Dam1 staple phosphorylation would promote Dam1c ring disassembly, explaining how kinetochore – microtubule attachments are destabilized and reset by the EC mechanism.

**One-Sentence Summary:** Phosphorylation disrupts the outer kinetochore to regulate kinetochore-microtubule attachments in mitotic error correction.

---

Chromosome segregation is an essential cellular function required for the equal propagation of genetic information from parent to daughter cells, achieved in eukaryotes through the kinetochore-mediated mechanical coupling of sister chromatids to the mitotic spindle (*1*). Kinetochores are large macromolecular assemblies delineated into the inner -chromatin binding-and outer -microtubule binding-complexes. The outer kinetochore couples the inner kinetochore centromere-binding CCAN (constitutive centromere associated network) to microtubules through three conserved complexes that together comprise the Knl1-MIND-Ndc80 (KMN) network (*2*). MIND interconnects CCAN with both the Ndc80 (Ndc80c) and Knl1 (Knl1c) complexes (*3-6*). Knl1c functions in the spindle-assembly checkpoint (*7*), whereas Ndc80c, a heterotetramer composed of the Ndc80, Nuf2, Spc24 and Spc25 subunits, is the major microtubule-binding component of the KMN network.

Outer kinetochore attachment to microtubules is augmented by additional essential microtubule-dependent kinetochore components. Many fungi, including *S. cerevisiae*, utilize the ten-subunit Dam1 complex, which self-assembles into a large ring around microtubules (*8-14*), whereas most metazoans contain the Ska complex (*2, 15-19*). Both the Dam1 and Ska complexes interact with Ndc80c to augment microtubule tip-tracking and rupture force *in vitro* (*20, 21*).

Although the Ndc80c and Dam1/Ska outer kinetochore complexes bind microtubules independently, full function and interaction strength requires both modules (*22-30*). A central unresolved question is how these complexes coordinately bind and track the dynamic microtubule plus-end to ensure kinetochores do not detach from the spindle. Moreover, as initial kinetochore-microtubule attachments are not necessarily bioriented, an error correction (EC) pathway, governed by antagonistic outer kinetochore phosphorylation by Aurora B kinase, ensures erroneous attachments, characterized by low tension, are weakened and reset (*31-37*). As kinetochores come under tension, a feature of successful biorientation, the outer kinetochore is dephosphorylated, attachments are stabilized, and the SAC is inactivated to trigger anaphase (*31-33, 35, 38-45*).

In yeast, the centromere-localized Aurora B kinase phosphorylates target sites in the Dam1c ring, either to suppress ring formation (*27, 29, 46*), or to block its association with Ndc80c (*22, 23, 42, 47*), resulting in weakened end-on attachment (*48, 49*). Phosphorylation of a second set of targets situated within the unstructured N-terminus of Ndc80 (Ndc80^N-Tail^), also contributes to EC. However, since this domain can be deleted without impairing cell viability (*26, 49, 50*), Ndc80^N-Tail^ phosphorylation appears subordinate to the role of Dam1c phosphorylation. Current models for error correction therefore propose that attachment reset occurs as a result of weakened outer kinetochore assembly, through prevention of Ndc80c-Dam1c contacts and suppression of ring assembly, and consequently reduced microtubule binding strength through phosphorylation of Ndc80^N-Tail^ and inhibition of co-operative binding. However, as there are currently no structures of a fully assembled outer kinetochore alone or in complex with the microtubule, the molecular mechanism by which EC phosphorylation antagonizes ring formation, and regulates kinetochore assembly at the Dam1c-Ndc80c interfaces remains fundamentally unknown.

To obtain a deeper understanding of the mechanism of outer kinetochore assembly on the microtubule and its regulation by error correction, we reconstituted and determined the cryo-EM structure of the budding yeast Ndc80c:Dam1c:microtubule complex. We observe multiple points of contact between Ndc80c and Dam1c, each of which contains embedded EC target sites, and the phosphorylation of which would be incompatible with complex assembly. An N-terminal ‘staple’ region of the Dam1 subunit assembles at the inter-protomer interface of the ring. The presence of embedded EC phosphorylation sites within the staple and Ndc80c-Dam1c interfaces indicates why EC would drive both Dam1c ring disassembly and destabilize kinetochore-microtubule attachments, respectively. We propose that EC thereby not only dismantles incorrectly attached outer kinetochore linkages, but also actively promotes attachment reset by driving turnover of phosphorylated Dam1. The re-establishment of kinetochore – microtubule attachments under low tension conditions, necessary for EC reset and sister chromatid biorientation, is achieved through unphosphorylated Dam1c rings tip-tracking shortening microtubules and rebinding kinetochores that have formed weak lateral-microtubule attachments. Consequently, these attachments convert to an end-on binding mode that can withstand the forces of chromosome segregation.

## Results

### Overall structure of the yeast outer kinetochore bound to microtubules

Outer kinetochore – microtubule complexes were assembled on cryo-EM grids through step-wise addition of taxol-stabilized microtubule filaments, Ndc80c and Dam1c. Cryo-electron micrographs show microtubules decorated with Dam1c rings and Ndc80c fibrils (**Fig. 1A**). In the consensus cryo-EM map of the outer kinetochore a prominent pattern of tandem decoration is observed in which Ndc80c coiled-coil ‘spokes’ emerge from the surface of the microtubule between diffuse Dam1c rings (**Fig. 1A, fig. S1A, B**). This decoration is semi-competitive or intermittent, insofar as the Ndc80 complexes appear not to occupy positions between the Dam1c ring and the microtubule lattice, and similarly Dam1c rings do not cap the Ndc80c spokes.

**Figure 1.**
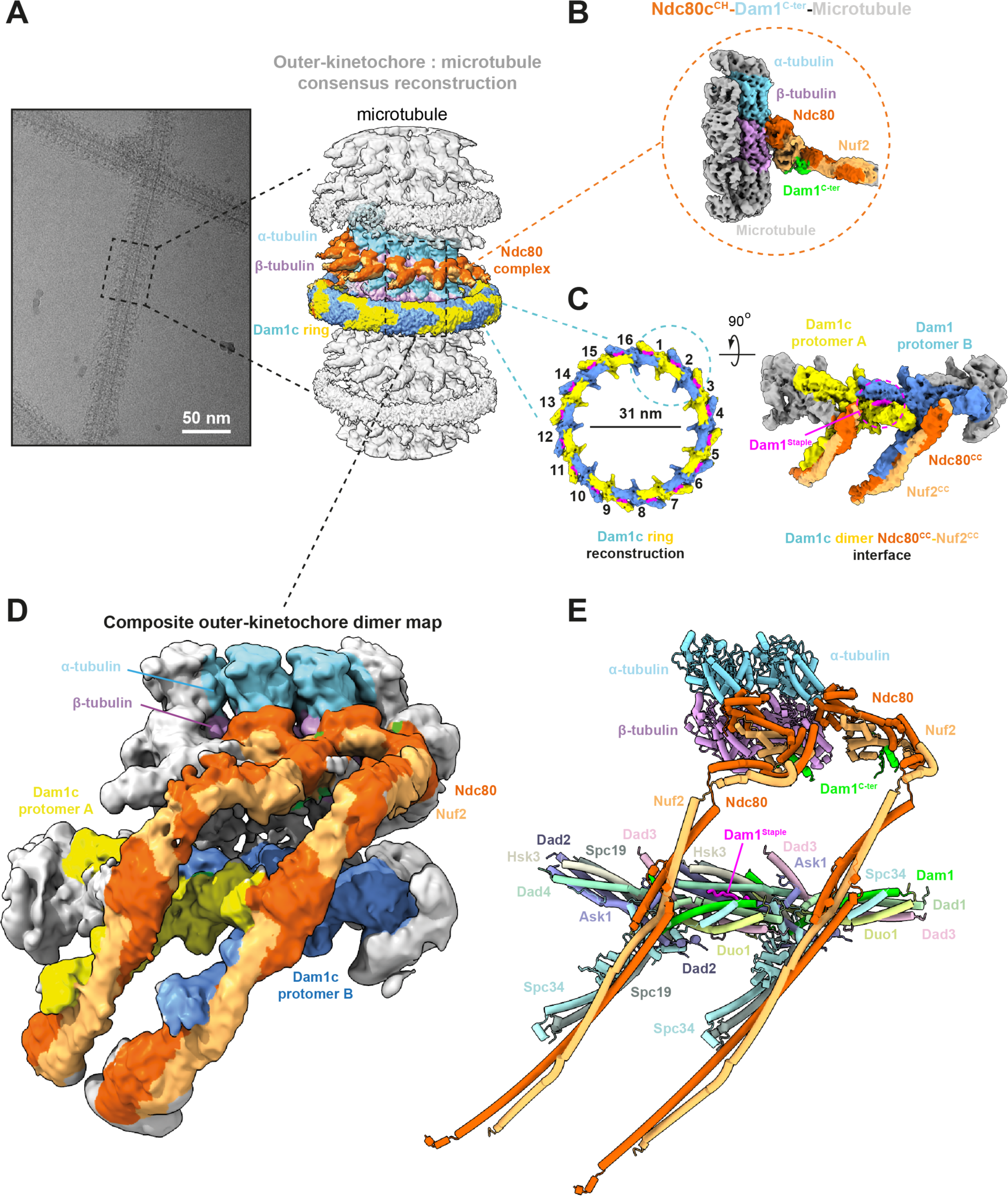
The yeast outer kinetochore complexes assemble through multiple interfaces to tandemly decorate the microtubule. **(A)** Representative cryo-EM micrograph and corresponding consensus reconstruction of the outer kinetochore microtubule complex. **(B)** Cryo-EM density map of the Ndc80c^CH^-Dam1^C-ter^-microtubule reconstruction. **(C)** Cryo-EM reconstruction of the 16-subunit Dam1c ring, and symmetry expanded Dam1c dimer Ndc80^CC^- Nuf2^CC^ interface. **(D)** Composite cryo-EM map and corresponding atomic model **(E)** of a yeast outer kinetochore microtubule dimer.

The intermediate resolution of the consensus cryo-EM map and the incompatible symmetries between the Dam1c ring, which we observed to have a copy number of 16 (as discussed below and consistent with (*12*)), and the 13 protofilament microtubule, necessitated a divide and consolidate strategy for structure determination (**Fig. 1B-D**, **fig. S1B, C, fig. S2A**). To determine a higher-resolution reconstruction of Ndc80c bound to the microtubule, we employed sub-boxing **(Materials and Methods)** (**fig. S2A, Table S1)**. In the resultant EM map, refined to an overall resolution of 3.54 Å, we observed additional density associated with Ndc80c that we attribute to the C-terminus of the Dam1 subunit (**Fig. 1B**), discussed below. In parallel, we performed signal subtraction around the Dam1c ring with a wide toroidal mask, which enabled us to obtain an intermediate reconstruction of the intact ring (**Fig. 1C, left panel**). By applying symmetry expansion to these particles, we could refine a dimer of Dam1c protomers to an overall resolution of 3.15 Å (**Fig. 1C, right panel, fig. S1C**). The coiled-coils of Ndc80c emanate from the calponin domains of the Ndc80 and Nuf2 subunits (Ndc80^CH^ and Nuf2^CH^) at the surface of the microtubule to fold across the outer surface of the Dam1c ring. Here we observed prominent density corresponding to the coiled-coil central region of the Ndc80-Nuf2 subunits (**Fig. 1C, D, fig. S1C**). Cryo-EM density is not visible for the Spc24-Spc25 subunits of Ndc80c. To obtain a complete model of the Ndc80-Nuf2:Dam1c:microtubule complex, we generated a composite map comprising two copies of the outer kinetochore protomer unit (defined as α/β tubulin:Ndc80c:Dam1c) by rigid body fitting of the separate volumes into the consensus outer kinetochore cryo-EM map (**Fig. 1D**). We then ‘folded’ the Ndc80-Nuf2 coils in the Dam1c^Protomer^ dimer component of the map until they were in proximity with the coils emanating from the Ndc80c^CH^ domains, and melded the models to generate a complete structure of the yeast outer kinetochore – microtubule complex (**Fig. 1D, E, Movie S1**).

This organization of the Dam1c ring and Ndc80c bound to the microtubule filament may explain how Dam1c augments outer kinetochore tracking of both polymerizing as well as depolymerizing microtubule tips (*23, 27, 28*). In the former scenario, Ndc80c is potentially simply borne along as a ‘passenger’ on the outer surface of the ring which need not necessarily form independent contacts with the microtubule, whereas in the latter, the complex is directly pushed through direct contacts with the ring, around which it folds as a hook. We anticipate this would result in ratcheting forward and re-binding of the Ndc80c calponin-homology domains to microtubules.

### Structure and regulation of Dam1c ring assembly by a Dam1 staple peptide

By performing particle image subtraction followed by focused classification and refinement without symmetry, we determined the structure of a 16-protomer Dam1c^Ring^ with an inner diameter of 31 nm (**Fig. 1C, fig. S1C, Table S1**). 2D classification without alignment of particles aligned on the ring showed that the Dam1c^Ring^ is tilted at variable angles relative to the microtubule (**fig. S1C, top left panel**), which presumably frustrated our attempts at obtaining a well-defined consensus outer kinetochore – microtubule reconstruction. Consistent with this observation, we find that the tubulin surface beneath the ring is unoccupied except for some diffuse density emerging from the α/β-tubulin C-terminal tails (**fig. S2B**), potentially representing the acidic E-hooks of α- and β-tubulin shown in some studies to enhance microtubule – Dam1c interactions (*9, 12*). Our structure confirms that the interaction between the Dam1c^Ring^ and microtubule potentially involves flexible participants, as also supported by biochemical and crosslinking mass spectrometry data (*11, 23, 46, 51*).

Dam1c^Ring^ assembly follows a shoulder-to-shoulder configuration in which the arms of each T-shaped Dam1c protomer form lateral contacts stacking above and below its clockwise and anti-clockwise neighbors, respectively (*11, 52*) (**Figs 1C**, **Fig. 2A**). To obtain higher-resolution information, we selected a single Dam1c^Protomer^ dimer position arbitrarily, reasoning that this would be the minimal informative repeat unit, then performed eight-fold symmetry expansion followed by iterative local refinement and subtraction (**Fig. 1C**, **Fig. 2A, fig. S1C**). Thereby, we derived a 3.2 Å map of a Dam1c^Protomer^ dimer, into which we placed and refined a structural model corresponding to two copies of the ten-subunit Dam1c (**Fig. 1E, 2A**). The inter-protomer junctions are similar to the cryo-EM structure of a truncated *C. thermophilum* Dam1c ring (*52*). In this structure, the map resolution was limited to 4.5 Å, and the model sequence was traced through reliance on well-defined bulky residues and validated by evolutionary direct-coupling analysis. In contrast, we find that in all ten subunits of *S. cerevisiae* Dam1c, most side-chains within the globular core of the complex are resolved, providing an unambiguous experimental model of the intact Dam1c^Protomer^ (**fig. S2C**).

**Figure 2.**
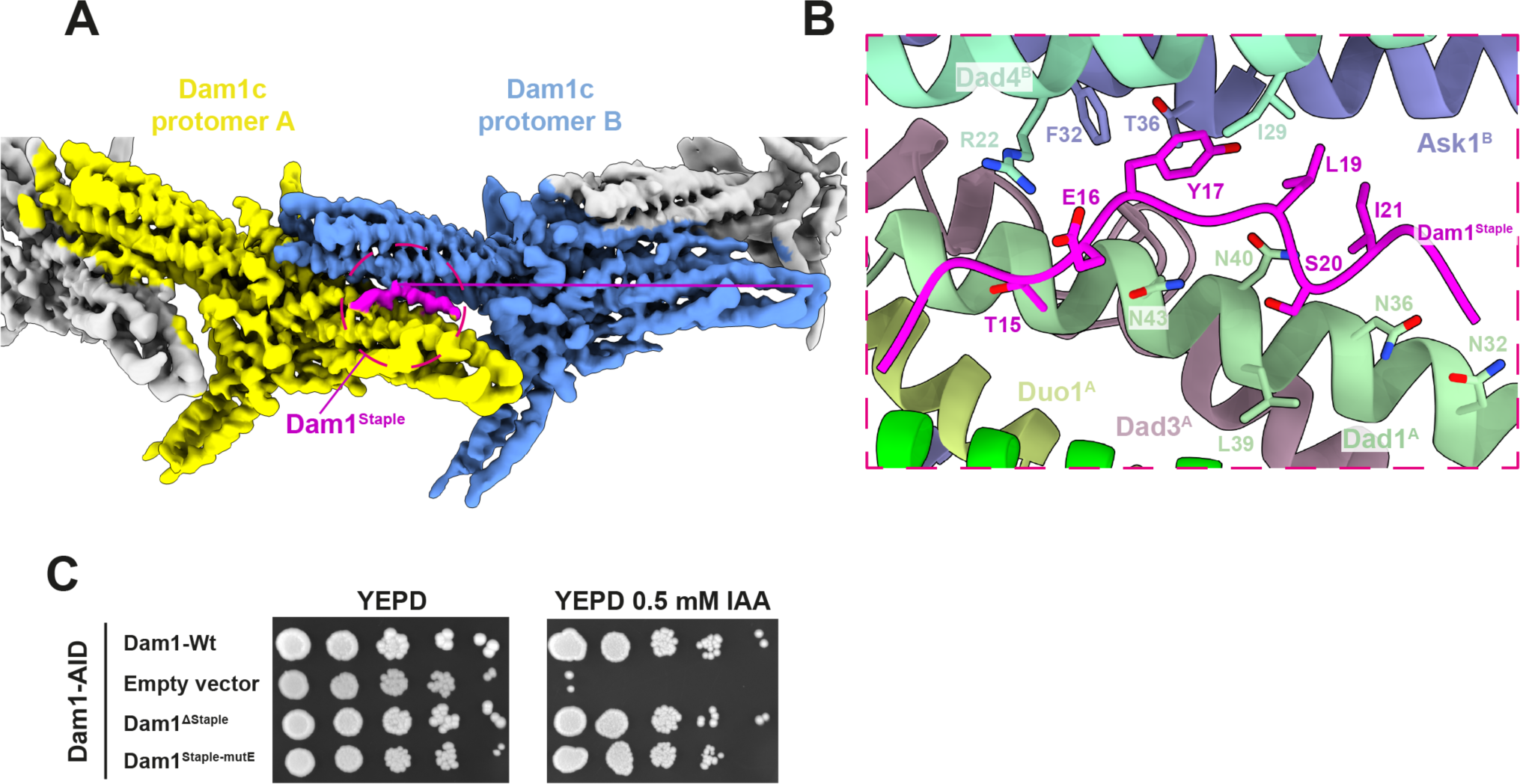
The Dam1 N-terminus forms a staple to stabilize Dam1c ring assembly that is negatively regulated by Aurora B kinase. **(A)** Structure of the Dam1c protomer dimer interface shows a staple density (purple) between protomer A (yellow) and protomer B (blue). (**B**) Details of amino-acid contacts at the Dam1c^Protomer^ dimer – Dam1^Staple^ interface. Ser20 is an Aurora B kinase phosphorylation site and is oriented toward the Dad1 subunit of protomer A. **(C)** Dam1 auxin depletion assays. Cells grown on YEPD agar and YEPD agar containing 0.5 mM IAA are shown side-by-side for each strain. Dam1^ΔStaple^: Dam1 mutant with residues 1-26 deleted, Dam1^Staple-mutE^: Dam1 mutant: Y17E/L19E/I21E.

Additionally, we resolved a ‘staple’ density located at the inter-Dam1^Protomer^ interfaces that was truncated from the *C. thermophilum* Dam1c expression construct (*52*). This staple (Dam1^Staple^) comprises the N-terminus (residues 1-26) of the Dam1 subunit of the Dam1c ring (**Fig. 2A, B**). The staple folds as a coil bridging the Dam1 and Dad1 subunits of Dam1c^ProtomerA^ with the Ask1 and Dad4 subunits of Dam1c^ProtomerB^. Contact with Dam1c^ProtomerA^ occurs mostly through hydrogen bonding between Dam1^Staple^ residues Thr15 and Ser20, and Dad1^A^ (**Fig. 2B**). Binding to Dam1c^ProtomerB^ is mediated by a salt bridge between Dad4^B^ Arg22 and Glu16 of Dam1^Staple^, a well as packing of Dam1^Staple^ residues Tyr17, Leu19, and Ile21 against a hydrophobic pocket generated by Ask1^B^ and Dad4^B^ (**Fig. 2B**). To test the requirement for this interface *in vivo*, we performed rescue assays in an *S. cerevisiae* strain with an auxin-inducible degron (AID) tag inserted at the C-terminus of the endogenous copy of Dam1 (**Table S2**). We then integrated a series of Dam1 variants lacking the AID tag at the *LEU* locus under control of their natural promoter and terminator. Cells grown on auxin with wild-type Dam1 as the ectopic copy rescued loss of endogenous Dam1, whereas cells carrying an empty vector failed to grow **(Fig. 2C**). To test the essentiality of the staple, we integrated two Dam1^Staple^ mutants into our Dam1-AID strain. These mutants comprise either deletion of the Dam1 staple (Dam1^ΔStaple^), or the combined mutation of Dam1 Tyr17, Leu19 and Ile21 to Glu (Dam1^Staple-mutE^). After plating these integrants on media containing auxin, we observed that the both the Dam1^ΔStaple^ and Dam1^Staple-mutE^ cells still could support viability in the absence of wild-type Dam1 (**Fig. 2C**).

Serine 20 of Dam1^Staple^ is a target of Aurora B kinase (*42*). In our structure, Ser20 is buried at the interprotomer interface, close to the main-chain of Dad1 of Dam1c^ProtomerA^, as well as Dam1c^ProtomerA^ residues Dad1 Leu39, Asn40 and Asn43 (**Fig. 2B**). Insertion of a phosphate at Ser20 would cause steric hindrance, as well as potential charge repulsion by backbone carbonyls, that would likely disrupt staple binding at this interface (**Fig. 2B**). Phosphorylation of Ser20 destabilizes Dam1c^Ring^ formation *in vitro* (*29, 46*). Similarly, mutations in the vicinity of the staple directly impair ring formation (*53*). However, these mutations can rescue loss of Aurora B kinase activity, presumably by driving increased kinetochore-microtubule attachment turnover (*53, 54*). Consistent with the notion that the Dam1^Staple^ regulates ring assembly, we found that the assembly of higher-order Dam1 complexes, as assessed by mass photometry, is substantially impaired when the staple is absent (**fig. S2D, E**).

### Structure of the Ndc80c:microtubule interface

Overall, the Ndc80c microtubule (MT)-binding domain forms a club-like structure comprising its two CH domains (**Figs 1A, E, Fig. 3A**). The underside of the club binds the α/β-tubulin interface along the lateral axis of the protofilament, whereas the coiled-coil shaft of Ndc80-Nuf2 projects outward at an angle approximately orthogonal to the CH domains. In contrast to the binding mode of humans and *C. elegans*, in which Ndc80c binds to every lateral α/β-tubulin interface (*55-57*), the yeast complex instead binds only at the α/β, but not the β/α site, resulting in intermittent binding along the microtubule protofilament (**fig. S3A**). Despite the differences in higher-order organization, the residues participating in the interaction between Ndc80c and α/β- tubulin are well-conserved between human and yeast, and are consistent with mutagenesis experiments showing the essentiality of interfacial amino acids in the Ndc80 subunit (*25, 26*). Consistent with its flexible structure, no EM density is visible for the Ndc80^N-tail^, and therefore we cannot account for how it may contribute to kinetochore – microtubule attachments.

**Figure 3.**
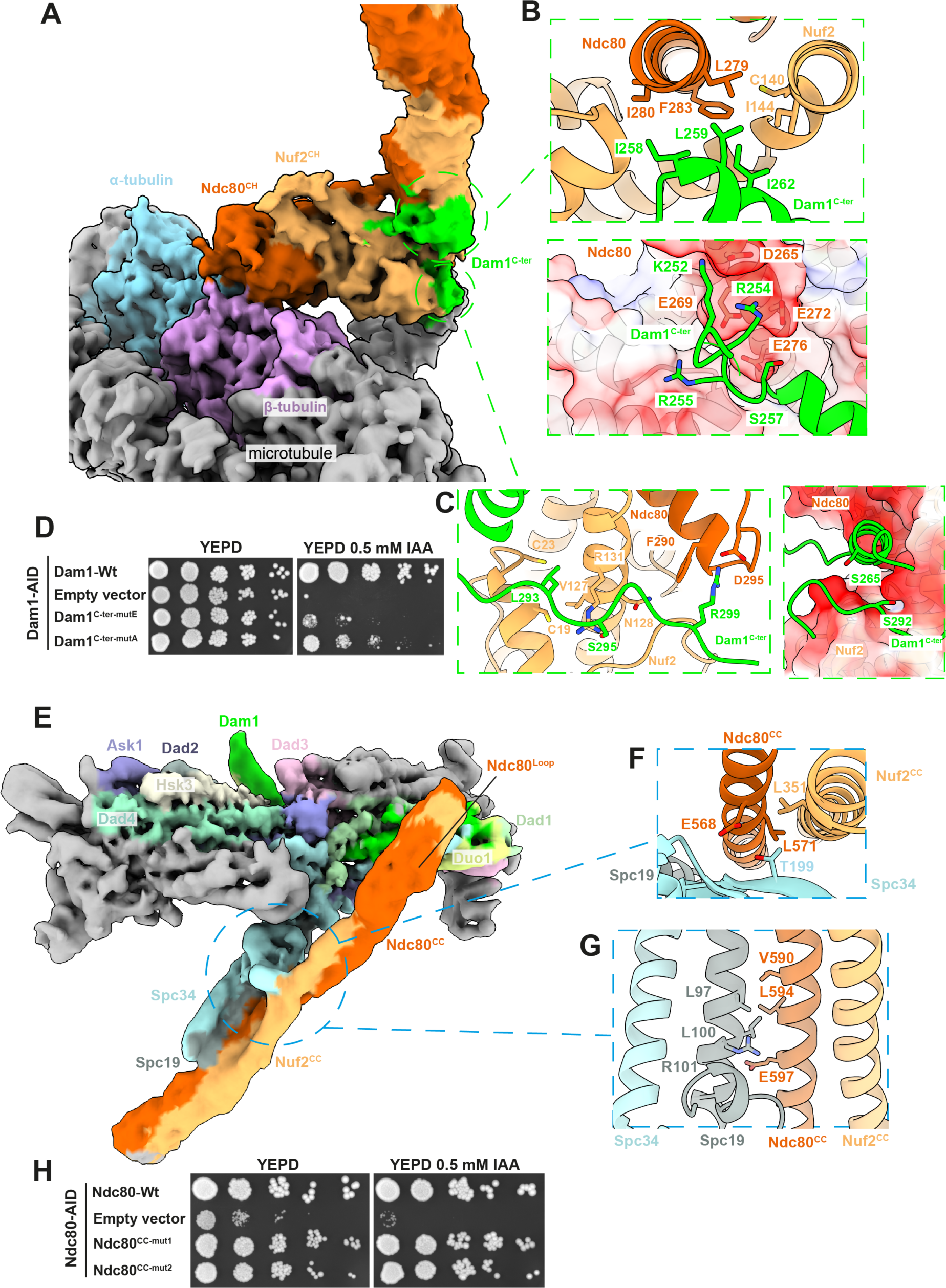
The Dam1 and Ndc80 complexes associate through a microtubule-proximal interface, and the outer surface of the Dam1c ring. **(A)** Cryo-EM reconstruction of the Ndc80c-Dam1c-microtubule interface. **(B)** The Dam1^C-ter^ α-helix packs against a hydrophobic interface generated by the Ndc80-Nuf2 coiled-coil emerging from the Ndc80c^CH^ domain (upper panel). Surface charges on Ndc80c^CH^ and position of the Ser257 error correction site on Dam1^C-ter^ are shown in the lower panel. **(C)** Structure of the Dam1^C-ter^ interface on the back-side of the Ndc80c^CH^ (left panel), positions of residues targeted by error correction and surface charges (right panel). Ser265 and Ser292 are oriented toward negative charges that would presumably drive dissociation of Dam1^C-ter^ from Ndc80^CH^ during EC. **(D)** Dam1 rescue assays on auxin. Cells grown on YEPD agar and YEPD agar containing 0.5 mM IAA are shown side-by-side for each strain. Dam1^C-ter-mutE^: Dam1 mutant: I258E/L259E/I262E, Dam1^C-ter-mutA^: Dam1 mutant: I258A/L259A/I262A **(E)** Cryo-EM reconstruction of the Dam1c monomer:Ndc80-Nuf2 coiled-coil interface. The position of the Ndc80^Loop^ is highlighted and fig. S3B. The Ndc80^CC^-Nuf2^CC^ binds Dam1c at two interfaces shown as insets: **(F)** Ndc80^CC^-Nuf2^CC^ dock against a β-strand on Spc34. The EC target site Thr199 is oriented toward Ndc80c. **(G)** The Spc19-Spc34 coiled coils pack against Ndc80^CC^-Nuf2^CC^ via residues in Spc19 and Ndc80. **(H)** Ndc80 auxin depletion assays. Cells grown on YEPD agar and YEPD agar containing 0.5 mM IAA are shown side-by-side for each strain. Ndc80^cc-mut1^: Ndc80 coiled-coil mutant1: E568A/V590A/L594A/E597A, Ndc80^cc-mut2^: Ndc80 coiled-coil mutant2: E568R/V590W/L594E/E597R.

### Binding of the Dam1 C-terminus to Ndc80c is essential and is regulated by error correction

We observed additional cryo-EM map density at the base of the Ndc80-Nuf2 coiled-coils emerging from the Ndc80c^CH^ domains that could not be accounted for by Ndc80c alone (**Fig. 3A**). While the additional density was not sufficiently well resolved to model *de* novo, crosslinking mass-spectrometry, and mutagenesis experiments suggest that this region of Ndc80c, termed the helical hairpin (**Fig. S3A**), interacts with Dam1c (*25*) by providing a binding site for the C-terminus of the Dam1 subunit of Dam1c (*23, 26*). To better understand this interface, we performed an AlphaFold2 run with Ndc80-Nuf2 together with residues 201 to 343 of Dam1. This predicted a tripartite complex with a configuration consistent with that reported in a subsequently published crystal structure (*58*). Two short segments from the C-terminus of Dam1 (Dam1^C-ter^) dock onto the Ndc80c^CH^ domains. One of these (residues 251 to 272) nestles in an amphipathic pocket at the base of the Ndc80-Nuf2 coiled-coils (**Fig. 3B**), and another (residues 287 to 301) snakes around the back-face of the club (**Fig. 3C**). A hydrophobic component of the Dam1^C-ter^:Ndc80c interface involves packing of Dam1 residues Ile258, Leu259, Ile262 against the CH domains of Ndc80-Nuf2 (**Fig. 3B, upper panel**). To interrogate the importance of this interface, we introduced Dam1 mutants bearing triple alanine or glutamate substitution of these residues as ectopic copies into our DAM1-AID strain (Ndc80^CC-mutA^ and Ndc80^CC-mutE^, respectively) (**Table S2**). Although these cells grew well on YEPD, upon IAA treatment growth is severely impaired, indicating that these mutants cannot fully substitute for wild-type Dam1 (**Fig. 3D**). Therefore, we conclude that the integrity of the Dam1^C-ter^:Ndc80c^CH^ interface is essential for proper kinetochore function and yeast cell viability.

During error correction, residues embedded at the Dam1^C-ter^:Ndc80c^CH^ interface are phosphorylated by Aurora B kinase, specifically Dam1 Ser257, Ser265, and Ser292 **(Figs 3B- lower panel, 3C-right panel)** (*41, 42*). Phosphorylation of these residues *in vitro* results in decreased strength and lifetime of outer kinetochore binding to the microtubule (*22, 23*), and phosphomimics compensate for loss of Aurora B kinase *in vivo* (*42*). To rationalize this, we mapped the surface charges of Ndc80c proximal to Dam1^C-ter^ in our model. Ser257 is oriented towards an α-helix on Ndc80c heavily decorated with negatively charged residues (**Fig. 3B, lower panel**). Phosphorylation of Ser257 would be sterically and electrostatically incompatible with binding of Dam1^C-ter^ to Ndc80c^CH^, thus weakening co-assembly of the outer kinetochore. Similarly, phosphorylated Ser265 and Ser292 would also be oriented toward negatively charged surfaces on Ndc80c (**Fig. 3C, right panel**).

### The central coiled-coil domain of Ndc80-Nuf2 folds across the outer rim of the Dam1 complex

In the consensus map of the outer kinetochore – microtubule complex, the trajectory of the Ndc80-Nuf2 coiled-coils brings them into proximity of the outer surface of the Dam1c^Ring^ (**Fig. 1B-D**). Consistent with this, we observed density in the cryo-EM maps of the Dam1c^Protomer^ dimer that could not be accounted for by the protein constituents of the Dam1c alone (**Fig. 1C, D, Fig. 3E**). This coiled-coil docks in close proximity to the inter-Dam1c^Protomer^ interface against a pair of β-strands from Spc34 that contain the EC target residue Thr199 (**Fig. 3E, F)**, and continues to run parallel with the coiled-coils comprising the C-terminal segments of Spc34 and Spc19 (**Fig. 3E, G**). Our structure is consistent with previous cross-linking mass spectrometry and insertion mutagenesis experiments indicating that the central segment of the Ndc80-Nuf2 coiled-coils proximal to the conserved Ndc80 loop directly interacts with the Spc34-Spc19 C-termini (*23, 25*). Following focused classification and flexible refinement, a distinct interruption in the coiled-coil became apparent (**Fig. 3E, fig. S3B**). This interruption corresponds to the Ndc80 loop region (Ndc80^Loop^), enabling us to define the amino-acid register of an AlphaFold2 prediction comprising the remaining coiled-coil. To lend further robustness to this interpretation, we mapped sequence conservation onto the structure, and observed that interfacial residues are generally well-conserved, whereas outward-facing residues are not (**fig. S3C**).

To test the impact of mutating this interface *in vivo*, we generated a yeast strain in which the endogenous copy of Ndc80 bore a C-terminal AID tag (**Table S2**). We then performed auxin depletion assays, and observed that whilst the introduction of an ectopic copy of wild-type Ndc80 rescued viability, the empty vector control could not (**Fig. 3H**). In contrast, mutation of a series of four conserved interfacial residues on Ndc80 did not impair viability, consistent with *in vitro* observations that disruption of this interface by phosphorylation of Spc34 at Thr199 exerts a very minor reduction in outer kinetochore – microtubule rupture forces (*22*). Indeed, our auxin-depletion experiments that disrupted the Dam1^C-ter^ and Ndc80c^CH^ interface (**Fig. 3D**) show that the Ndc80c coiled-coil:Spc34 interface is not sufficient for proper outer kinetochore function (*25, 26*).

### Disruption of outer kinetochore interfaces weakens microtubule attachments *in vitro*

The full load-bearing potential of the outer kinetochore and its ability to track dynamic microtubule ends is dependent on co-assembly of Ndc80c and Dam1c (*22-30, 47*). Disruption of coordinated binding to the microtubule is a central proposed mechanism for how error correction regulates attachments (*22-25, 27, 28, 47–49, 59*).

The forces imposed by the yeast spindle on the kinetochore during metaphase have been estimated to be in the vicinity of ∼8-16 pN per kinetochore – microtubule attachment (*60*). Correspondingly, *in vitro* optical trap experiments, in which purified yeast kinetochores and reconstituted kinetochore – microtubule attachments are challenged to determine the force at which they rupture, have established values in the order of ∼5-10 pN (*22, 24, 27, 61*). To test the contribution each of the interfaces presented in our structures make to the ability of the outer kinetochore to withstand force, we purified a series of Dam1 variants and performed force-rupture assays. In these assays, adapted from (*22, 62*), Ndc80c was immobilized on streptavidin-coated beads via a SNAP-biotin tag and beads held in the optical trap laser were then brought into close proximity of dynamic microtubules immobilized on glass cover slides, permitted to bind and then withdrawn to determine at what force the attachments ruptured. We determined a median rupture force of 5.4 pN for Ndc80c alone that increased to 10.4 pN in the presence of Dam1c (**Fig. 4A**), confirming that the complete Ndc80c:Dam1c outer kinetochore complex withstands greater forces than does isolated Ndc80c (*22, 24, 27*). By contrast, addition of variants including mutation of the Dam1^C-ter^ (Ndc80c^CH^ contact-1) and Dam1^Staple^ (ring assembly contact) substantially lowers the outer kinetochore rupture forces to 5.5 pN and 6.5 pN, respectively, whereas the Ndc80^CC^ (Spc19-Spc34:Ndc80-Nuf2 coiled-coil contact-2) essentially has no effect on rupture strength when combined with Dam1, after accounting for reduced basal microtubule binding of this Ndc80 mutant (**Fig. 4A, B**). Hence, mutation of outer kinetochore assembly through the Dam1^C-ter^- Ndc80c^CH^ interface or disruption of Dam1 ring formation can independently weaken microtubule binding. It was previously reported that introduction of either phosphonull or phosphomimics at both the Dam1^Staple^ and Dam1^C-ter^ severely impairs viability of yeast cells (*42*). Cumulatively, our structural and biochemical results suggest that ablation of these functional elements of Dam1, targeted for inhibition by EC, would generate complexes that are unable to assemble rings, contact Ndc80, nor to bear load (**Fig. 4A, B**). To test whether concurrent disruption of these elements of Dam1, would impact kinetochore function *in vivo*, we introduced combined Dam1^Staple^ and Dam1^C-ter^ mutants into our Dam1-AID strain (**Table S2**). We observed that mutagenesis of key Dam^C-ter^ residues to alanine (Dam1^C-ter-mutA^) permitted weak growth that was completely abolished when the Dam1^Staple^ was deleted or mutated (Dam1^ΔStaple^ and Dam1^Stable-mutE^, respectively) (**Fig 4C**).

**Figure 4.**
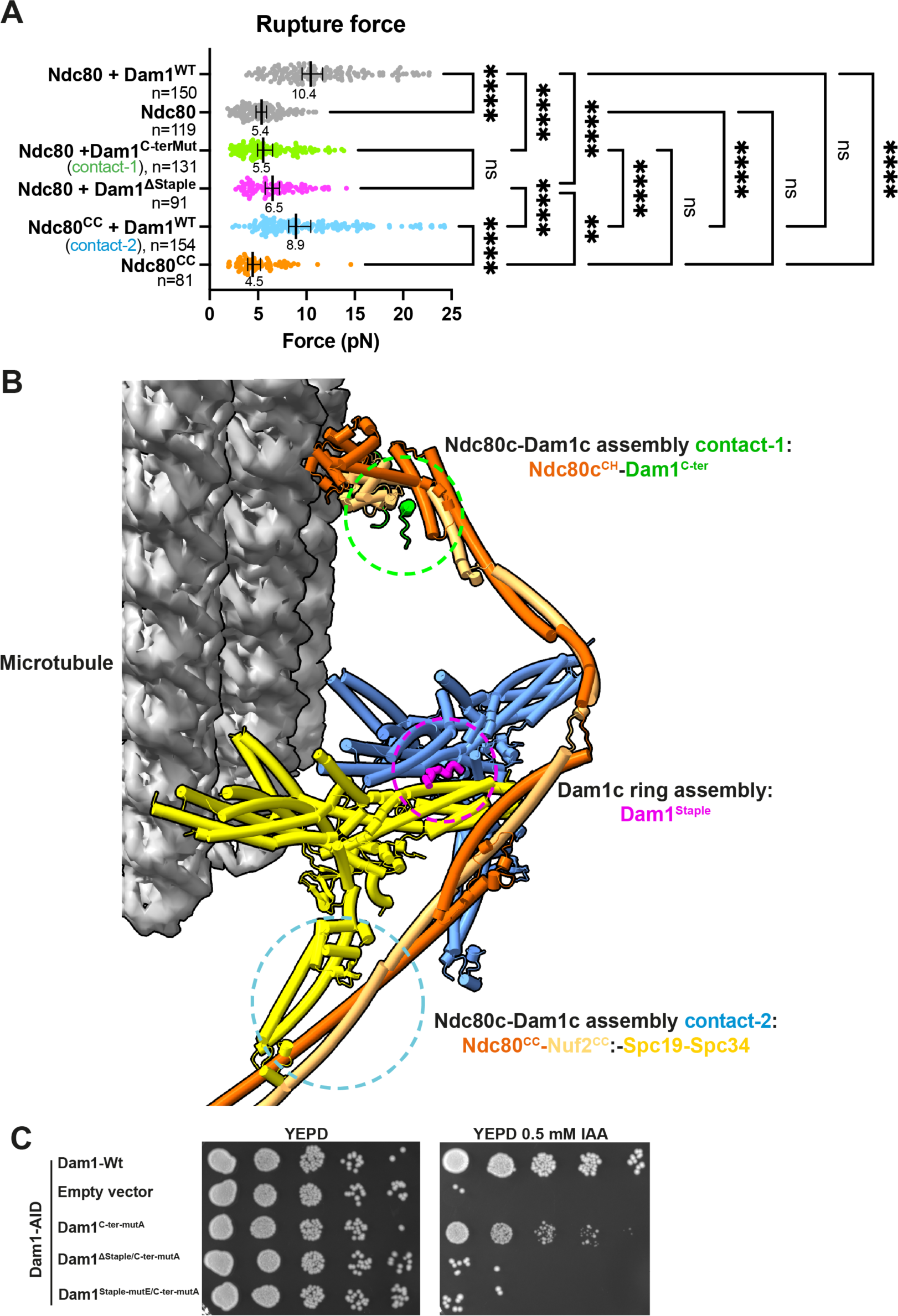
A structural model for mitotic error correction and chromosome segregation by the yeast kinetochore. **(A)** Force-rupture measurements of outer kinetochore – microtubule complexes. Each circle represents a single rupture event (the maximum trap force before rupturing). The total number of measurements for each condition are indicated by n values. The black bar represents the median rupture forces, with 95% CIs. Number below the black bar indicates median values. A Kruskal-Wallis test to determine which medians are significantly different was performed, ****: p<0.0001, **: p= 0.0046, ns: not significant. **(B)** Structure of an outer kinetochore dimer bound to the microtubule. Outer kinetochore and Dam1c ring assembly contacts that are targeted by error correction are highlighted. **(C)** Dam1 auxin depletion assays. Cells grown on YEPD agar and YEPD agar containing 0.5 mM IAA are shown side-by-side for each strain. Dam1^C-ter-mutA^: Dam1 mutant: I258A/L259A/I262A, Dam1^Staple-mutE^: Dam1 mutant: Y17E/L19E/I21E. Simultaneous mutation of the Dam1 Staple (Dam1^Staple-mutE^) and Dam1 C-terminus (Dam1^C-ter-mutA^) results in lethality.

Consistent with the notion that EC directly drives attachment turnover by breaking outer kinetochore assembly, our force-rupture data correspond well to measurements of *in vitro* reconstituted Dam1 complexes phosphorylated at the respective interfaces by Aurora B (Ndc80c^CH^ and Ndc80-Nuf2 coiled-coil contacts) (*22, 24, 27*), as well as for complexes containing a phospho-mimetic Dam1 S20D mutation (Dam1^Staple^ ring assembly contact) (*24, 48*). Finally, the severity of the force-rupture phenotypes measured here correlates closely with the degree of viability defect caused by the corresponding mutants in our auxin depletion assays. Hence, mutagenesis of the interfaces identified in our structures likely causes reduced fitness *in vivo* because these cells form defective kinetochore – microtubule attachments that cannot support normal chromosome segregation.

## Discussion

Kinetochore-mediated chromosome segregation is an essential facet of eukaryotic life. Here we consolidate previous structural findings together with *in vivo* measurements of relative kinetochore subunit positions and stoichiometry with our structure of the outer kinetochore to generate a structural model for a metaphase yeast holo-kinetochore bound to the microtubule plus end (**Fig. 5A, B**) (*3, 63-66*). Prior structural observations have shown that the incorporation of the yeast outer kinetochore into CCAN is mediated by three distinct pathways (*3, 66-69*). In this framework, four copies of MIND bind two molecules of CENP-U embedded within CCAN and two molecules of CENP-U recruited via the CENP-A N-terminus (*66*). A further two copies of MIND are separately recruited by CENP-C. These six MIND complexes then bind six Ndc80c. Finally, the two CENP-T subunits of CCAN directly recruit an additional two Ndc80c, giving rise to eight Ndc80c outer kinetochore linkages to the microtubule. The eight Ndc80c molecules are then readily accommodated by the 16 binding sites provided by the assembled Dam1c ring, which is recruited through Ndc80c in a microtubule-dependent fashion (*70*). The excess of Dam1c-binding sites over Ndc80c also provides for the avidity necessary for the mechanism of biased diffusion that enables Ndc80c to processively track dynamic microtubule ends (discussed below) (*71*).

**Figure 5.**
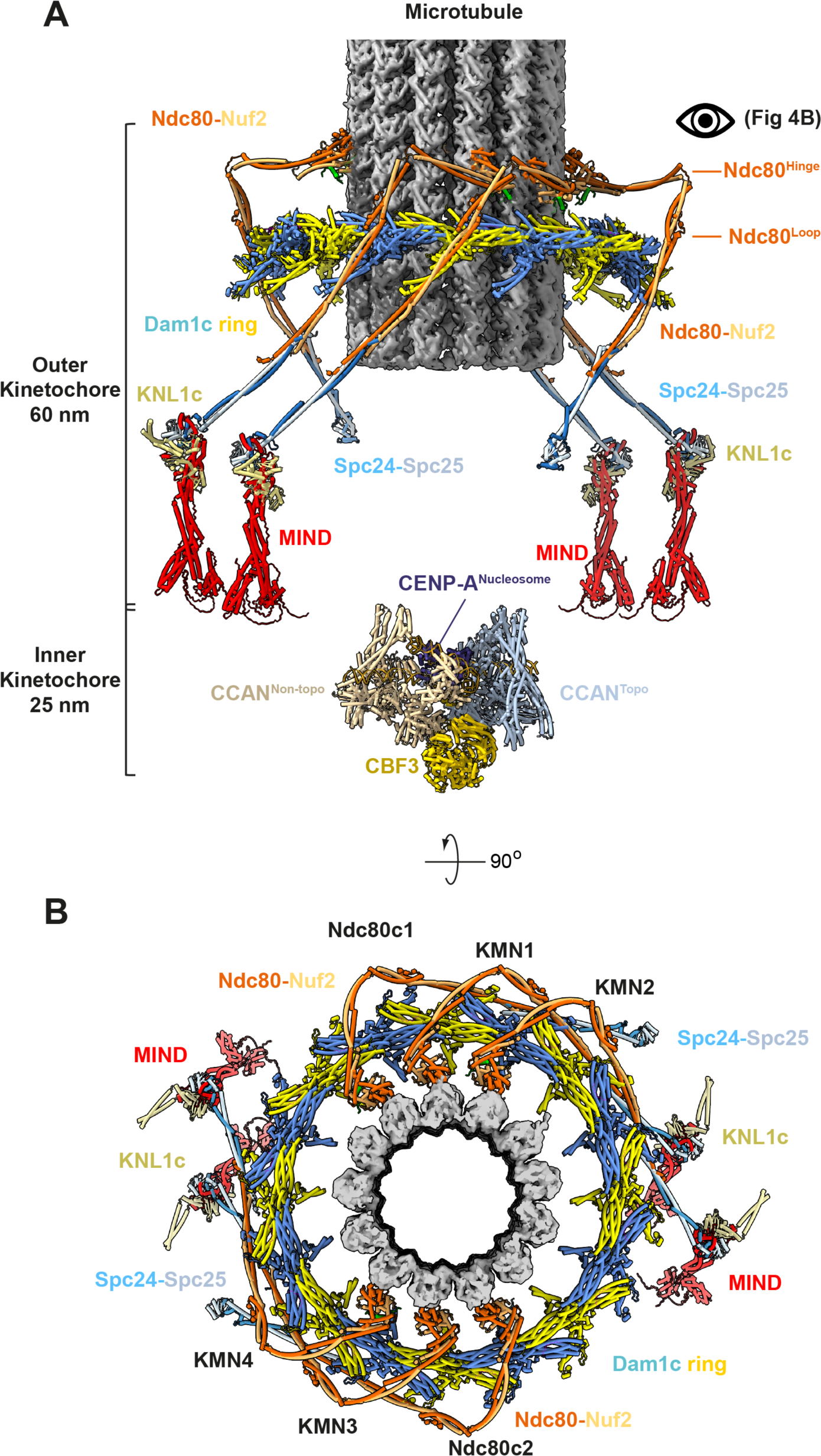
A structural model for mitotic error correction and chromosome segregation by the yeast kinetochore. **(A)** Structural model of the complete yeast kinetochore bound to a microtubule. Shown are the microtubule, Dam1c ring, four KMN network complexes, two Ndc80 complexes and the CCAN inner kinetochore complex (comprising two CCAN promoters, CBF3 and CENP-A nucleosome). The flexible linkers connecting CCAN to the KMN network and Ndc80c are not shown. The view is in plane with the microtubule helical axis. **(B)** Top view of a complete outer kinetochore – microtubule complex viewed from the microtubule minus-end.

We observe a configuration of individual Ndc80c molecules which, mediated by a flexible hinge (*72*), have folded as hooks across the Dam1c ring surface. This explains the ∼37 nm separation between the N-terminus of Nuf2 and C-terminus of Spc24, observed for budding yeast kinetochores at metaphase using high-resolution colocalization methods (*64*) that is 19 nm shorter than expected for the fully extended Ndc80c (*72, 73*). The hinge between Ndc80c^CH^ and Ndc80^Loop^ permits a degree of flexure (*72*) that explains how the coiled-coils of Ndc80-Nuf2 can dock in a regular parallel configuration across Spc19-Spc34 of Dam1c despite adherence of the Ndc80c^CH^ to the pseudo-helical rise of the microtubule surface (*64, 74*). This may have important implications for translocation, as it means that variable positioning of Ndc80c on the microtubule can be accommodated without inducing strain that would break its connection with the Dam1c ring, and would permit Ndc80c to swivel forward and rebind during microtubule depolymerization toward the spindle pole during chromosome segregation.

The central function of EC is to promote bioriented kinetochore – microtubule attachments. The mechanism by which errors are detected is an area of active investigation, however several proposed models have gained prominence, in which as kinetochores come under tension, the targets of EC are less effectively phosphorylated by a centromere-localized pool of Aurora B kinase (discussed in (*33, 75*)). Consistent with this, in yeast cells tension inversely correlates with outer kinetochore phosphorylation (*45*). Observations that ectopic targeting of Aurora B kinase to kinetochores that have bioriented is sufficient to detach them from the spindle in human (*76*) and yeast cells (*77*) are consistent with the spatial separation model. In the alternative, but not incompatible conformational change model, structural alterations within kinetochores under tension render them resistant to phosphorylation (*78*). Shortening of yeast, *Drosophila* and human kinetochores as tension is reduced is indicative of structural changes that are consistent with both models (*63*).

Our metaphase yeast holo-kinetochore model (**Fig. 5A, B**) suggests how kinetochore components are organized along the kinetochore-microtubule axis. We positioned the inner kinetochore just below the MIND complex. In this configuration, the end-to-end dimension of the kinetochore is ∼85 nm, a value consistent with the relative location and separation of kinetochore components measured in metaphase yeast cells using fluorescence localization microscopy (*64, 65*). Shortening of the kinetochore as tension is reduced (*64*) could be facilitated through the flexible linkers connecting the inner kinetochore with MIND and Ndc80 complexes.

Yeast error correction is trifurcated between disruption of Dam1c ring assembly, disassembly of inter-Dam1c:Ndc80c interfaces that weaken cooperative assembly on the microtubule, and the phosphorylation of Ndc80^N-Tail^ (*22-24, 26–28, 41, 42, 47*). Collectively these inhibitory activities have the dual consequence of dismantling the outer kinetochore and dissolving kinetochore – microtubule attachments, however the precise molecular mechanisms by which EC achieves this turnover have remained elusive.

We found that outer kinetochore complex assembly is mediated through multiple interfaces that are directly antagonized by Aurora B kinase phosphorylation, whereas Dam1c ring assembly is facilitated by a newly discovered ‘staple’ that physically bridges Dam1c protomers and contains an embedded EC target. In contrast to human Ndc80c, we do not observe inter-Ndc80c contacts, nor can we account for the flexible Ndc80^N-Tail^ (*55, 79*). Although yeast cells with either deletion of Ndc80^N-Tail^ (*28, 49, 80, 81*), or Ndc80^N-Tail^ EC-site mutations (*50, 81*) are viable, Ndc80^N-Tail^ nevertheless contributes to EC. Disruption of EC via a phospho-null Ndc80^N-Tail^ mutant is synthetic lethal with a ts mutant of Aurora B kinase (*50*). *In vitro*, Ndc80^N-Tail^ contributes to microtubule binding (*26, 82*) and phosphomimics of Ndc80^N-Tail^-EC sites weaken kinetochore-microtubule attachments (*48*), consistent with the inability of yeast cells with an Ndc80^N-Tail^ deletion to generate proper tension between sister chromatids (*81*). Cell biological and *in vitro* experiments showed that the Dam1, Ask1 and Spc34 subunits of Dam1c form a network of interactions between the Dam1c ring and Ndc80c that promote cooperative outer kinetochore assembly on the microtubule and are subject to disruption by EC (*22, 23, 25, 26*). The region implicated in Ask1:Ndc80c binding is too mobile to resolve in our cryo-EM maps, but is most likely situated at the Ndc80c hinge between the Ndc80c^CH^ and Ndc80c loop (*23*). Our molecular models therefore directly account for two EC-sensitive contacts: the Dam1^C-ter^:Ndc80c^CH^ and the Spc34:Ndc80-Nuf2 coiled-coil interfaces. At both interfaces, residues phosphorylated in EC would cause electrostatic and steric repulsion (**Fig. 2B**, **Fig. 3B, C**). Our structural model therefore provides insight into the mechanism by which EC-mediated phosphorylation directly suppresses complete outer kinetochore assembly.

A fundamental paradox at the heart of EC is how initially weak, low-tension kinetochore attachments can then escape re-phosphorylation during attachment reset to form full-affinity connections to the microtubule that are able to withstand tension. Ndc80c retains a degree of microtubule affinity even when Ndc80^N-tail^ is either deleted (*26, 82*) or incorporates EC phosphosites (*47*), therefore enabling Ndc80c to form microtubule attachments when tension is absent. However, it has previously been unclear how Dam1c could be reincorporated into the outer kinetochore if the Ndc80c:Dam1c interfaces are abolished during EC. The major targets of phosphorylation that govern cooperative outer kinetochore assembly are concentrated within the Dam1 complex (*22, 23, 41, 42*). Thus, achieving full attachment strength would either require dephosphorylation and/or replacement of Dam1c. Phosphorylation of Dam1 at Ser20 blocks the formation of higher-order Dam1 complexes and increases its diffusion on microtubules (*24, 27*). In our structure, we discovered that this EC phosphorylation site targets a staple that bridges adjacent Dam1c protomers, providing a molecular explanation for how EC directly controls ring assembly (**Fig. 2**). We therefore propose that simple replacement of Dam1c by EC-mediated turnover would resolve the EC-reset paradox by exchanging phosphorylated Dam1c, that cannot interact with Ndc80c, for non-phosphorylated Dam1c. Following EC phosphorylation, kinetochores remain attached to the lateral face of the microtubule (*49, 83*), and the phosphorylated Dam1c diffuses away. However, Dam1c not associated with the kinetochore remains unphosphorylated (*43*) and is transported to the kinetochore via independent tracking of the microtubule plus-ends. Upon encountering the kinetochore, unphosphorylated Dam1c is able to associate with Ndc80c to facilitate conversion from lateral to end-on attachment (*47, 49, 83*). This Ndc80c:Dam1c assembly, even with Ndc80^N-Tail^ phosphorylated, would be sufficient to drive displacement of the outer kinetochore and associated conformational change, rendering it resistant to the centromere-localized Aurora B kinase (*69, 84*). Consequently, protein phosphatases would remove inhibitory phosphorylation from Ndc80^N-Tail^, generating full-strength kinetochore – microtubule attachments. Our findings therefore provide mechanistic insight into how the combined effects of weakened kinetochore – microtubule attachments and dissolution of the Dam1c ring not only disrupts incorrect attachments but also facilitates their reassembly by promoting reincorporation of non-phosphorylated Dam1c to generate full-affinity outer kinetochore linkages to the microtubule.

Although microtubule-associated motor proteins play facilitating roles in chromosome congression and biorientation, the central engine of chromosome motility is the microtubule itself (*83, 85, 86*). Hydrolysis of GTP by tubulin generates bending strain that is released as protofilaments curl outward from the microtubule during disassembly (*87, 88*). Two prominent models posit how this mechanical energy is exploited by the kinetochore to segregate chromosomes: the conformational wave, and biased diffusion. In the former, the curling protofilaments directly lever the kinetochore toward the spindle poles (*85*). Additionally, as the Dam1c ring has a narrower diameter than the disassembling protofilaments, it would serve as a cuff to track the microtubule (*8, 10, 11, 62*). In the biased diffusion model, an array of diffusive kinetochore attachments detaches and rebinds the tubulin lattice as it depolymerizes, thereby tracking the microtubule (*89*). A key prerequisite of this model is multivalency, satisfied by the multimerization of the Dam1c ring, as well as the recruitment of multiple Ndc80 complexes, up to eight, by a single yeast inner centromere complex (*65, 66*). Our structural findings provide support for and reconcile both proposals. The presence of multiple docking sites for Ndc80c within Dam1c indicates that the ring complex can fulfill the role of a topological coupler to receive pulling forces exerted by the curling protofilaments and directly generates multivalency by coordinating multiple Ndc80 complexes to enable biased diffusion. Constraining Ndc80c through relatively flexible linkages permits ratcheting of Ndc80c whilst presumably favoring rebinding to the same microtubule, consistent with observations that Dam1c confers processivity to the kinetochore-microtubule attachments (*10, 26–28, 30, 47, 49*). In addition to buried peptidic and coiled-coil interfaces, the folding of the Ndc80c hook around the outer circumference of the Dam1c suggests a topological mechanism by which the ring facilitates tracking of depolymerizing microtubules by Ndc80c. The intermittent decoration of the microtubule that we observe implies that Dam1c may not only facilitate Ndc80c translocation through a tethering mechanism, but might actively drive its displacement during microtubule depolymerization.

Our findings support a model for kinetochore-mediated chromosome segregation wherein the ring acts as a topological sleeve that is pushed along by the curved protofilaments of the depolymerizing microtubule, and drives Ndc80c translocation through steric occlusion. Association of the complexes through flexible peptidic interfaces effectively mediates a tethering mechanism which prevents Ndc80c from detaching completely from the microtubule when displaced by the advancing Dam1c ring, instead allowing it to ratchet forward and rebind, thus pulling sister chromatids toward the spindle poles.

The forces applied on the centromere are minimal in anaphase, thus we anticipate that our structure reflects the likely state of a kinetochore as it performs chromosome segregation. However, future work will be required to address if and how this structure is modified by the application of tension by the mitotic spindle during chromosome alignment and biorientation.

## Acknowledgments

We acknowledge Diamond Light Source for access and support of the cryo-EM facilities at the UK’s national Electron Bio-imaging Centre (eBIC) [under proposal EM BI31336], funded by the Wellcome Trust, MRC and BBRSC. We are grateful to the LMB EM facility for help with the EM data collection, to J. Grimmett, T. Darling and I. Clayson for high-performance computing, and to J. Shi for help with insect cell expression and Vicente Jose Planelles Herrero for help with the optical tweezer study. We thank Stan Yatskevich for discussions and Noah Turner for comments on the manuscript. For the purpose of open access, the author has applied a CC BY public copyright license to any Author Accepted Manuscript version arising.

## Funding

UKRI/Medical Research Council MC_UP_1201/6 (DB)

Cancer Research UK C576/A14109 (DB)

## Author contributions

Conceptualization: D.B. and K.W.M.

Methodology: KWM, CB, TD, JY, ZZ, AB, DB

Investigation: KWM, CB, TD, JY, ZZ, Visualization: KWM

Funding acquisition: DB

Project administration: DB, KWM

Writing – original draft: KWM

Writing – review & editing: KWM, DB, CB, TD, JY, ZZ, AB

## Data and materials availability

All data are available in the main text or the supplementary materials. PDB models and cryo-EM maps are available through the RCSB and EMDB. Accession numbers are provided in Table S1.

## Supplementary Materials

Materials and Methods

Figs. S1 to S4

Tables S1 to S2

References (90-108)

Movies S1

