## Supplementary Materials for "Mechanism of outer kinetochore assembly on microtubules and its regulation by mitotic error correction"

**The PDF file includes:**

Materials and Methods
Figs. S1 to S4
Tables S1 to S2
References 90-108

**Other Supplementary Materials for this manuscript include the following:**

Movies S1

### Materials and Methods

Cloning of the Dam1 and Ndc80 complexes

The ten genes encoding the yeast Dam1 complex (Dam1c) comprising Ask1, Hsk3, Dad1, Duo1, Dad2, Spc19, Dad3, Spc34, Dad4, Dam1, and the four genes encoding the Ndc80 complex, comprising Spc25, Ndc80, Spc24 and Nuf2 were amplified by PCR from *Saccharomyces cerevisiae* genomic DNA (strain s288c) and cloned into the pU1 plasmid, using a modified Multibac expression system as described previously (*90*). Double Strep-II tags preceded by a TEV cleavage site were inserted at the C-termini of Dam1 and Ndc80. Ask1, Hsk3, Dad1, Duo1, Dad2, Spc19 were initially cloned into pU1, then further co-assembled into a pU2 vector. The pU1 expression cassettes individually containing Spc19, Dad3, Spc34, Dad4 were co-assembled into a pF2 vector. For the Ndc80 complex, pU1 expression cassettes containing Spc25, Ndc80, Spc24 and Nuf2 were assembled into a pF2 vector. A single virus for expressing the Dam1 complex was generated from bacmids of the combined pU2 and pF2 vectors, and a virus for Ndc80 complex was made from its pF2 vector as described (*90*).

Expression and purification of the Dam1 and Ndc80 complexes

A single baculovirus for expressing the Dam1 complex was generated from bacmids of the combined pU2 and pF2 vectors, and a baculovirus for Ndc80 complex was made from its pF2 vector as described (*90*). The Dam1 and Ndc80 complexes were expressed separately in High Five insect cells, which was not tested for mycoplasma contamination and was not authenticated. For Dam1c and Ndc80c, two liters each of High Five cells were infected with 2.5% v/v of P3 cell culture, and the cells were harvested 48-72 hours post-infection by centrifugation (2000xg; Beckman JLA 8.1).

For Dam1c and its associated mutants, High Five cells were lysed by sonication in Dam1c lysis buffer (50 mM Tris pH8.0, 150 mM NaCl, 1 mM EDTA, 1 mM DTT), supplemented with 8 mM benzamidine-HCl, EDTA-free protease inhibitors (Roche), and benzonase (Invitrogen). Lysate was then clarified by centrifugation and applied to a Strep-Tactin column (Qiagen) coupled to an ÄKTA Pure. Protein immobilized on the resin was then washed with 10 column volumes of lysis buffer, and eluted with lysis buffer containing 5 mM desthiobiotin (Sigma-Aldrich). Dam1c eluate was then supplemented with 10 mM ATP, 10 mM $MgCl_2$, 20 mM KCl, and incubated overnight at 4 °C. Subsequently, the eluate was diluted to an NaCl concentration of 50 mM, loaded on to a Resource Q anion exchange column (Cytiva), and washed with 10 column volumes of ATP wash buffer (50mM NaCl, 20 mM Tris pH8.0, 1 mM EDTA, 1mM DTT, 10 mM ATP, 10 mM $MgCl_2$, 20 mM KCl). Dam1c was then eluted over a salt gradient from 50 mM to 450 mM NaCl. Peak fractions containing the protein complex were then pooled, concentrated (Centricon, Amicon), and applied to a Superdex S200 column (Cytiva) equilibrated in a buffer containing 25 mM HEPES pH 7.4, 150 mM NaCl pH7.4, 1 mM EDTA, 1 mM DTT. Peak fractions containing Dam1c were then harvested, concentrated to 2.1 mg/ml, flash frozen in liquid $N_2$, and stored at -80 °C as 100 µL aliquots.

For the Ndc80 complex and its associated mutants, cell pellets were lysed by sonication in Ndc80c lysis buffer (50mM Tris pH8.0, 200 mM NaCl, 8% glycerol, 0.5mM EDTA, 2 mM DTT, 2mM benzamidine, 0.2mM PMSF), supplemented with EDTA-free protease inhibitors (Roche) and benzonase (Invitrogen). After centrifugation the Ndc80 complex was purified by affinity chromatography (Strep-Tactin (Qiagen)) in a buffer containing 50 mM Tris-HCl pH 8.0, 250 mM NaCl, 0.5 mM EDTA and 2mM DTT, followed by elution with 5 mM desthiobiotin, and removal of the Double StrepII tags by TEV protease. The Ndc80 complex was further purified by anion exchange chromatography (Resource Q (Cytiva)) in a starting buffer of 20 mM Tris-HCl pH 7.5, 100 mM NaCl, 1 mM EDTA and 2 mM DTT, followed by elution using an NaCl gradient from 100 mM to 1 M. Fractions containing the Ndc80 complex were then pooled, concentrated and applied to a Superdex S200 size-exclusion chromatography column (Cytiva) equilibrated in a buffer containing 20 mM Tris-HCl pH 7.5, 200 mM NaCl, 1 mM TCEP. Peak fractions containing the Ndc80 complex were then concentrated to 5 mg/mL, flash frozen in liquid $N_2$ and stored at -80 °C as 50 µL aliquots.

<u>Microtubule polymerization</u>
Lyophilized porcine tubulin was purchased (Cytoskeleton, Inc.) and resuspended in polymerization buffer (70 mM NaCl, 25 mM MES pH 6.5, 1 mM EGTA, 1 mM EDTA, 3 mM $MgCl_2$, 3 mM GTP) to a concentration of 10 mg/ml, flash frozen in liquid $N_2$ and stored at - 80 °C as 10 µl single-use aliquots. To polymerize microtubules, thawed tubulin stocks were diluted 1:1 with warm polymerization buffer and incubated for 20 minutes at 37 °C. The polymerization mixture was then mixed 1:1 with warm polymerization buffer containing 40 µM taxol (Cytoskeleton, Inc.), followed by centrifugation at 12,000 r.c.f. for 8 minutes in a bench-top centrifuge (Eppendorf), after which the supernatant was discarded and the microtubule pellet resuspended by vigorous pipetting in 200 µl BRB80-IT (80 mM PIPES pH 6.8, 1 mM EDTA, 1 mM EGTA, 1 mM DTT, 20 µM Taxol and 0.05% IGEPAL-CA 630). Microtubules intended for vitrification on cryo-electron microscopy (cryo-EM) grids were then diluted 20fold in BRB80-IT and kept at 22 °C.

<u>Reconstitution and vitrification of the outer kinetochore – microtubule complex for cryo-EM</u>
Purified Dam1c and Ndc80 complexes were dialyzed for 1 hour at 4 °C into BRB80 buffer (80 mM PIPES pH 6.8, 1 mM EDTA, 1 mM EGTA, 1 mM DTT) using 200 µL capacity Slide-A-Lyzer capsules (ThermoFisherScientific) with a molecular weight cut-off of 20 kDa. The protein complexes were then centrifuged in a bench-top centrifuge for 10 minutes at a speed of 20,000 r.c.f., diluted to a working concentration of 2.5 µM in BRB80 supplemented with 0.05% IGEPAL-CA 630 and kept at 22 °C.

For the assembly of outer kinetochore – microtubule complexes, 2 µL of the diluted microtubule stock was first applied to glow-discharged Quantifoil 300 mesh gold R2/2 grids (Quantifoil Micro Tools), and inserted into a Vitrobot IV device operating at a temperature and humidity of 30 °C and 100%, respectively. The microtubules were then allowed to adsorb to the grids for 1 min, prior to application of 2 µL of the Ndc80c stock (2.5 µM), and further incubated for 1 min. Finally, 2 µL of the Dam1c stock was added (2.5 µM), and the entire mixture left to incubate for a period of 3 minutes prior to blotting (blot time 1s, blot force -15) and plunge-freezing in liquid ethane maintained at a temperature of 93 K using a home-built cryostat device (*91*).

<u>Cryo-EM data acquisition</u>
Micrograph movies were collected with a Titan Krios microscope operating at 300 keV, using a K3 camera (Gatan) and energy filter (20 eV), at a magnification of 81k, yielding a nominal pixel size of 1.06 Å. In total 39.61k micrograph movies were recorded at a dose rate of 16.6 $e^-$/px/s

with 3s exposure and 50 frames. Automated data-collection was performed in EPU (ThermoFisherScientific), with aberration-free image-shift. Defocus values ranged from -1.2 to -3.0 µm with an interval of 0.2 µm.

Cryo-EM data processing

Motion correction and contrast-transfer function estimation were performed in RELION 4.0 (*92*) using MotionCor2 (*93*) and CTFFIND (*94*), respectively. An initial set of outer kinetochore – microtubule particles were picked manually from a randomly selected set of 300 micrographs, to provide input coordinates to train a filament-picking model in crYOLO (*95*). All micrographs were then picked using crYOLO in filament mode, with a spacing of 81 Å. Resulting start-end coordinates were then imported into RELION 4.0 and used to extract corresponding particle images from the original micrographs, with 4x binning. Particles were then subject to two rounds of 2D classification, using a helical rise of 82 Å, to exclude junk, and over-lapping or broken microtubule particles. Subsequently, the Microtubule RELION-based Pipeline (MiRP) (*96*) was used to classify the microtubules according to protofilament (pf) number. The distribution of pf numbers across the dataset was: 11pf (4.7%), 12pf (40.7%), 13pf (42.7%), 14pf (6.3%), 15pf (3%), 16pf (2.6%). As 13 pf microtubules are the predominant species observed in yeast mitotic spindles, we selected only this fraction for further analysis. At this stage, particles were exported to cryoSPARC (*97*), and subjected to several rounds of heterogeneous refinement. In the initial round, the particles were classified against 6 identical reference models corresponding to a 13 pf MT low-pass filtered to a resolution of 20 Å. Limiting the maximum alignment resolution to 30 Å was essential to prevent the microtubule itself from dominating alignment, which otherwise resulted in smearing out of the Dam1c ring map density. In subsequent rounds, maps containing the most consistent Ndc80 decoration and continuous Dam1c rings were iteratively fed back through heterogeneous refinement until arriving at a set of output volumes which appeared to differ only in the relative register of the Ndc80c-Dam1c on the microtubules. At this stage, particles were exported back into RELION 4.0, and for each set 3D consensus refinement was performed. In order to consolidate the particles into the same register, a synthetic single-turn 13pf ring was manually placed using Chimera into equivalent positions in each of the (four) volumes, resampled onto the coordinate space of the target volume using vop, and the center-of-mass determined using the relion_image_handler --com command, followed by particle re-extraction, with 4x binning and the relevant x,y, and z shifts required to harmonize the particle register. All particle sets were then joined and subjected to another round of 3D consensus refinement in RELION, followed by 3D classification without alignment (T=8) to exclude any residual mis-aligned, poorly decorated, or junk particles. All particles were then extracted without binning, re-refined, and subjected to two rounds of CTF refinement followed by Bayesian polishing, and a final round of consensus 3D refinement. Thereafter, the procedures used for particle classification diverged into several distinct streams:

*Stream 1. The Ndc80c-Dam1$^{C\text{-}ter}$-microtubule focused reconstruction.*

In essence, to determine a higher-resolution reconstruction of Ndc80c bound to the microtubule, we employed sub-boxing **(fig. S2A),** in which we centered a coordinate on each Ndc80c:α/β-tubulin interface in the consensus map, and re-extracted this expanded particle set for further processing. As described above, refinement of decorated microtubules yielded a reconstruction centered on the microtubule lumen in which the microtubule was oriented along the z-axis and both the Ndc80c-microtubule interface and Dam1c rings were visible at intermediate resolution.

We leveraged the multiplicity of the Ndc80c-microtubule interface in our data to improve the reconstruction in this region. The positions of visible Ndc80c-microtubule interfaces in the consensus reconstruction were annotated in napari (*98*). A consistent local coordinate system was defined programmatically for each Ndc80c-microtubule interface in which the new z-axis was perpendicular to the microtubule and the new y-axis was oriented along the protofilament in the +z direction of the consensus reconstruction. A rigid body 3D transformation, composed of a shift and a rotation, was calculated from these data and used to recenter and reorient particles from the consensus reconstruction on each Ndc80c-microtubule interface was then calculated, and applied to the consensus particle set. This analysis was implemented in Python using the packages starfile, NumPy and einops. Subsequent 3D reconstruction of particles extracted (with 2x binning) at these positions with their associated orientations yielded a reconstruction centered on the Ndc80c-microtubule interface in the expected orientation, confirming the correct transformation had been applied and providing an initial reference for subsequent refinement.

After an initial round of 3D consensus refinement, to improve the homogeneity of the particle set a mask encapsulating a single Ndc80c-Dam1$^{C\text{-}ter}$- α/β-tubulin was applied during 3D classification without alignment (T=20, classes=4) in order to separate particles according to Ndc80c occupancy. After excluding particles which were either poorly aligned or lacked Ndc80c, a second round of 3D classification without alignment (T=24, classes=4) using a mask only around the Ndc80 density was pursued. Particles were then unbinned, and subjected to a single round of 3D refinement with a mask around a single Ndc80c-Dam1$^{C\text{-}ter}$, but including rows of flanking α/β-tubulin. Particle subtraction was then performed in RELION using a mask focused on a single Ndc80c$^{CH}$-Dam1$^{C\text{-}ter}$ with neighboring α/β-tubulin. Thereafter, particles were imported into cryoSPARC, where two iterations of local refinement, with a round of particle subtraction in between, were performed yielding a map of the Ndc80c-Dam1$^{C\text{-}ter}$-microtubule complex at an overall resolution of 3.2 Å.

*Stream 2. Dam1 complex focused reconstruction.*
To obtain a focused reconstruction of the Dam1c ring, a toroidal mask was generated using a previously determined model of the Dam1c ring complex filtered to a resolution of 200 Å, which was then used for particle subtraction around the central ring of the outer kinetochore – microtubule consensus map. Ring particles were subsequently refined against an input model of a 16-protomer Dam1c ring, which was previously shown to be the dominant copy number obtained *in vitro* around microtubules. At this stage, no symmetry was applied. Particles were then classified in 2D with alignment in order to remove particles for which residual microtubule remained. A further round of refinement, with symmetry relaxation to C16, resulted in a ring consensus map with a resolution of 5.3 Å. To validate the presumptive copy number, parallel analysis of the particles was performed using 3DVA in cryoSPARC and 3D classification without alignment. In both cases, the principle source of variability was not copy number, but partial fragmentation of the ring, as well as modest doming. Therefore, to obtain a higher resolution consensus volume for the subtracted ring, a final round of refinement was performed in RELION with C16 symmetry applied, resulting in a final symmetrized ring reconstruction at a higher overall resolution of 5 Å.

*Stream 3. The Dam1c protomer-dimer staple focused reconstruction.*

To obtain higher resolution reconstructions of a fundamental repeating unit of the Dam1c ring, the cleaned ring particle set following the 3D consensus refinement step with symmetry relaxation above was exported into cryoSPARC, and subjected to homogeneous refinement with C16 symmetry applied. The particle set was then expanded by applying C8 symmetry, and subjected to local refinement using a mask and model corresponding a Dam1c protomer dimer. Particle subtraction was then performed, and the particles sorted using 3D classification, in cryoSPARC. After discarding less well-defined particles, a final round of local refinement was performed yielding a volume containing a Dam1c protomer dimer with apparent density for the Dam1 N-terminal staple, to an overall resolution of 3.15 Å.

*Stream 4. The Dam1c dimer Ndc80-Nuf2 central coiled-coil focused reconstruction.*
The same Dam1c input set used in stream 3 was retained in RELION and subjected to 3D refinement with C16 symmetry, followed by C16 symmetry expansion, and re-refinement. From the resulting 3D volume, a prominent albeit less well-defined coiled-coil density emanating from the outer surface of the ring could be observed and could not be attributed to the Dam1 complex alone. To improve the resolution of this coiled-coil, particles were subtracted using a generous mask including the Dam1c protomer and coiled-coil, then subjected to 3D refinement. The resulting particle sets were then classified in 3D without alignment (T=20, 8 classes), classes bearing more defined coiled-coils were pooled iterative 3D classification and 3D refinement. Particles were then imported into cryoSPARC, and subjected to a round of local refinement followed by 3DVA (*99*). Particles contributing to the most continuous variability volume were then locally refined and subjected to 3DFlex-EM (*100*) using default parameters. In the resulting maps, the coiled-coils of Ndc80-Nuf2 and Spc34-Spc19 were clearly resolved. The Ndc80-Nuf2 coiled-coil register could be assigned due to unambiguous density corresponding to the Ndc80 loop region, which forms a winding 'switch-back' motif distinct from the neighboring coiled-coils.

*Stream 5. The outer kinetochore – microtubule complex consensus volume with an aligned seam.*
The particles arising from 3D refinement of the consensus outer kinetochore – microtubule reconstruction were subjected to 3D classification without alignment in RELION (T=4, 6 classes), leading to a single class with an apparently well-defined seam as determined from direct inspection of the inner surface of the microtubule. A further round of 3D refinement in RELION resulted in a reconstruction of 4.24 Å (e.g. Nyquist for the 2x binned particles) in which heterotypic packing of neighboring α/β-tubulin was clearly apparent at the seam location.
Stream 6. Dam1 complex focused reconstruction from seam-aligned consensus.
Particle subtraction was performed as described in stream 2 and a single round of 3D refinement was performed in RELION with symmetry relaxation to C16, resulting in a reconstruction of the Dam1c ring essentially identical to that derived from the larger non-seam aligned dataset.

<u>Cryo-EM model building and refinement</u>
For all models, geometry and fit-to-map was optimized via real-space refinement was in PHENIX (*101*).

*Stream 1. The Ndc80c-Dam1$^{C\text{-}ter}$-microtubule complex.*
A previously determined structure of porcine α/β-tubulin (with bound taxol, GTP, and GDP), and an Alphafold2 prediction of the Ndc80-Nuf2-Dam1$^{C\text{-}ter}$ were fit as rigid bodies into the

corresponding map using Chimera (*102*). Side-chains were apparent for the α/β-tubulin, as well as the neighboring contacts with Ndc80, and these regions were therefore manually remodeled into the density in Coot (*103*). Density for the base of the calponin-homology domains of Ndc80-Nuf2, where Dam1$^{C\text{-ter}}$ is bound, was less well defined, and this region as well as the adjacent coiled-coils of Ndc80-Nuf2 were therefore modelled more conservatively using Isolde (*104*) as implemented in Chimera X (*105*). Weak density for the α/β-tubulin tails was apparent but could not be unambiguously modelled, and similarly the linker connecting the two segments of the Dam1 C-terminus could not be observed in the map and is thus omitted.

*Stream 3. The Dam1c protomer-dimer staple complex.*
Two copies of a decameric Dam1 complex, predicted by Alphafold2 were manually placed and rigid body fitted into the map with Chimera. The model was then manually corrected in Coot, with flexible regions being excised. Once all globular domains had been accounted for, the map was visually inspected for additional unassigned densities, which revealed a short stretch of amino acids that were manually sequenced from the map, and that correspond to the Dam1 N-terminal staple. As the linker region connecting the staple to the globular segment of Dam1 spans 55 amino acids, this peptide was assigned to the Dam1 chain of protomer 1, however we cannot exclude that it may originate instead from its neighbor.

*Stream 4. The Dam1c dimer Ndc80-Nuf2 central coil complex.*
The model of the Dam1c protomer dimer staple complex derived from stream 3 was manually fit in the map together with the Ndc80-Nuf2 subunits of an AlphaFold2 prediction of the complete Ndc80 complex, and adjusted with rigid body fitting in Chimera. The register of the Ndc80-Nuf2 coiled-coils could be placed unambiguously according to the position of the Ndc80 loop domain in the density maps. Due to the lower local resolution of the Spc34-Spc19-Ndc80-Nuf2 interface, this region was re-modelled conservatively using restrained flexible fitting in Isolde.
A complete Ndc80c-Dam1c-α/β-tubulin dimer complex was modelled by aligning the Dam1c dimer Ndc80-Nuf2 central coil map onto the outer kinetochore – microtubule consensus map, with the fitmap command in ChimeraX. The separate models corresponding to the Dam1c dimer Ndc80-Nuf2 central coil complex, and two copies of the Ndc80c-Dam1$^{C\text{-ter}}$-α/β-tubulin complex were placed by rigid body fitting into the former and latter maps respectively. The coiled-coil regions of Ndc80-Nuf2 were then separately re-modelled using restrained flexible fitting in Isolde, until a short stretch of residues that over-lapped between the separate models were considered to be in close proximity. Thereafter, the Dam1c dimer Ndc80-Nuf2 central coil, and the (Ndc80c-Dam1$^{C\text{-ter}}$-α/β-tubulin)2 models were exported separately on the origin of the outer kinetochore consensus maps, and the Ndc80-Nuf2 chains consolidated manually in Coot. To maintain reasonable geometry at the Ndc80-Nuf2 hinge domain, where the coiled-coils fold abruptly in order to dock on the outer surface of the Dam1c ring, the complete model of the dimeric outer kinetochore - α/β-tubulin complex was subject to a further round of restrained flexible fitting in Isolde.

<u>AlphaFold2 predictions</u>
AlphaFold2 (*106*) predictions were run on locally on the MRC LMB cluster, with default parameters, including a relaxation step. For the Dam1 complex, all 10 subunits were compiled into a single FASTA document and submitted. For the Ndc80 complex, the four subunits were

compiled as above into a single FASTA file and submitted. For the Ndc80c-Nuf2-Dam1$^{C\text{-}ter}$ complex, full-length Ndc80-Nuf2 and residues 201 to 243 of Dam1 were submitted together.

Yeast strains

Yeast strains are listed in Table S2. To construct Dam1-mAID and Ndc80-mAID the mAID insert was amplified from pST1933 (*107*) using Q5 polymerase using primers containing 48bp 5' and 3' extensions bearing homology to the corresponding loci in the yeast genome.
Yeast cells from exponential culture were harvested, washed with sterile dH2O and vortexed with transformation buffer (200 mM lithium acetate, 40% PEG 3350, 50 µg salmon sperm DNA (Invitrogen, 15632011) together with the transformation cassettes. Cells were heat shocked for 45 minutes at 42 °C, then allowed to recover in YEPD for 2 hours at 30 °C. Transformants were then plated on YEPD Agar containing 500 µg/ml G418 (Sigma, A1720-5G) and grown at 30 °C for 3 days. Colonies were screened by PCR using primers targeted to the 3' of the target gene and an internal region within the KanMX cassette. PCR products of positive transformants were then sequenced to confirm in-frame integration of the mAID tags at the appropriate site.

Auxin depletion assays

10 ml cultures of each strain were grown overnight in YEPD at 30 °C. Cells were then diluted to an OD600nm of 0.1 and serially diluted 1:10 over 5 dilutions. 4 µl of each titration was then plated on agar plates containing YEPD (for the negative control experiment) and YEPD supplemented with 0.5 mM auxin (Sigma, cat. No. I3750-100G-A). Cells were then grown at 30 °C for 3 days and imaged.

Optical trap rupture force assay

All rupture force experiments were performed on a LUMICKS C-Trap Edge setup. Biotinylated Ndc80 complex was linked to 0.9 µm-diameter streptavidin-coated polystyrene beads (Lumicks). 3.5 pM beads were incubated with 30 nM Ndc80c for 1 h at 22 °C and washed twice in PBS.

Flow cells were assembled essentially as previously described (*108*). Briefly, glass 22x22 mm coverslips (NEXTERION, Schott) were incubated for at least 48 h at 22 °C in a 1:10 (w/w) solution of mPEG-Silane (30 kDa, PSB-2014, Creative PEGWorks) and Biotin-PEG-Silane (3.4kDa, Laysan Bio Inc.) in 96 % (v/v) ethanol and 0.2 % (v/v) HCl. The coverslips were then washed in ethanol and ultrapure water, dried with an air duster and assembled into a flow cell on a slide using double-sided tape (Adhesive Research AR-90880 precisely cut with a Graphtec CE6000 cutting plotter). The chamber was first perfused with one chamber volume of BRB80 solution (80mM PIPES pH 6.9, 2 mM $MgCl_2$, 0.5 mM EGTA, 1mM DTT). Then with one chamber volume of 5% Pluronic-F127 in BRB80 and immediately flushed out with 8 chamber volumes of BRB80. One chamber volume of Neutravidin (25mg/ml) diluted in BRB80 was left to incubate for 5 min at 22 °C and then flushed out with 4 chamber volumes of BRB80. Biotin-GMPCPP stabilized microtubule seeds (20% HiLyte 488-tubulin, 20% biotin-tubulin 60% tubulin (Cytoskeleton, Inc.) were then injected and let to adhere to the neutravidin for 5 minutes. The chamber was then washed with 10 chamber volumes of BRB80. Ndc80c coated beads were added to the reaction mixture (1x BRB80, 16.7 µM of Tubulin, 80 µM GTP, 1mg/ml BSA) Dam1 complex or the Dam1 mutants were added to the reaction mix to achieve a final concentration of 30 nM as required. The microscope stage was kept at 28ºC with an objective and condenser heater (Lumicks). 488nm HiLyte Tubulin seeds were briefly localized in TIRF mode. Microtubule growth, bead capture and rupture experiments were carried out using IRM (Interference reflection microscopy). Data were collected for a maximum of 90 minutes after addition of the reaction mixture.

Rupture-force experiments were based on published procedures (*22, 62*). Beads were held in the laser trap approximately 200 nm above the surface and the position was continually monitored at 40 KHz. Trap stiffness was calibrated using a power spectrum method and was in the range of 0.025-0.035 $pN.nm^{-1}$. Beads were allowed to bind to the matrix near to the tip of the microtubule. The laser-trap was moved at 10 $\mu m.s^{-1}$ until the bead ruptured from the tip. Data were recorded using Lumicks software and rupture forces were manually determined by examining force and position versus time traces using PRISM. 95% confidence intervals (CIs) were generated for the median rupture forces. Normality test showed a non-parametric analysis should be performed therefore statistical significance for the difference in medians between each sample was determined using a Kruskal-Wallis test.

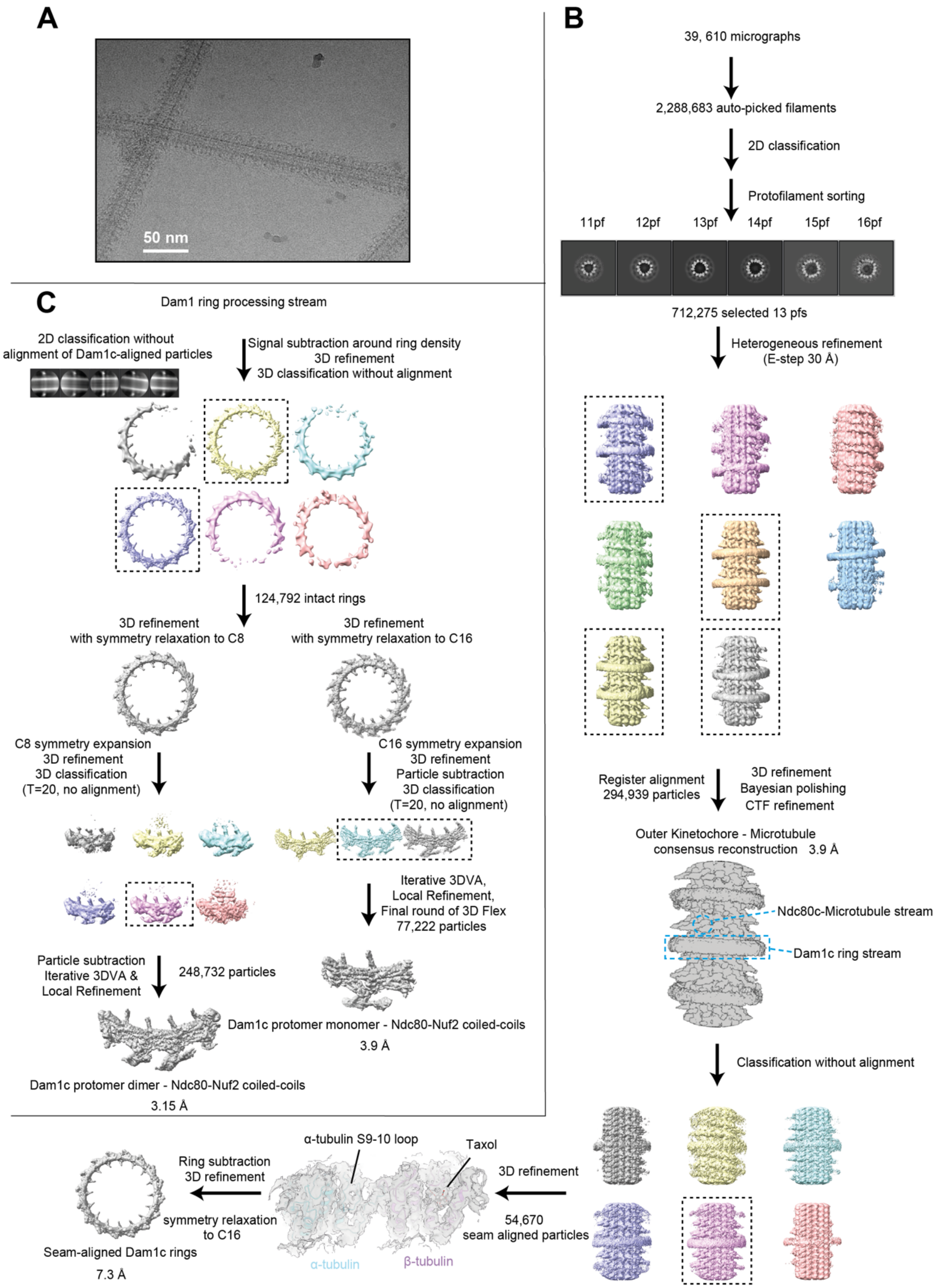


**Figure S1. Workflow for the consensus and Dam1c ring focused outer kinetochore – microtubule complex cryo-EM reconstructions. (A)** Representative cryo-EM micrograph of the outer kinetochore microtubule complex. **(B)** Workflow for cryo-EM processing of the consensus outer kinetochore model and seam-aligned Dam1c ring reconstruction. Map density and models corresponding to the microtubule seam interface is shown. **(C)** Workflow for the Dam1c ring-focused cryo-EM processing stream. Representative 2D classes showing variable ring orientation relative to the microtubule are shown.

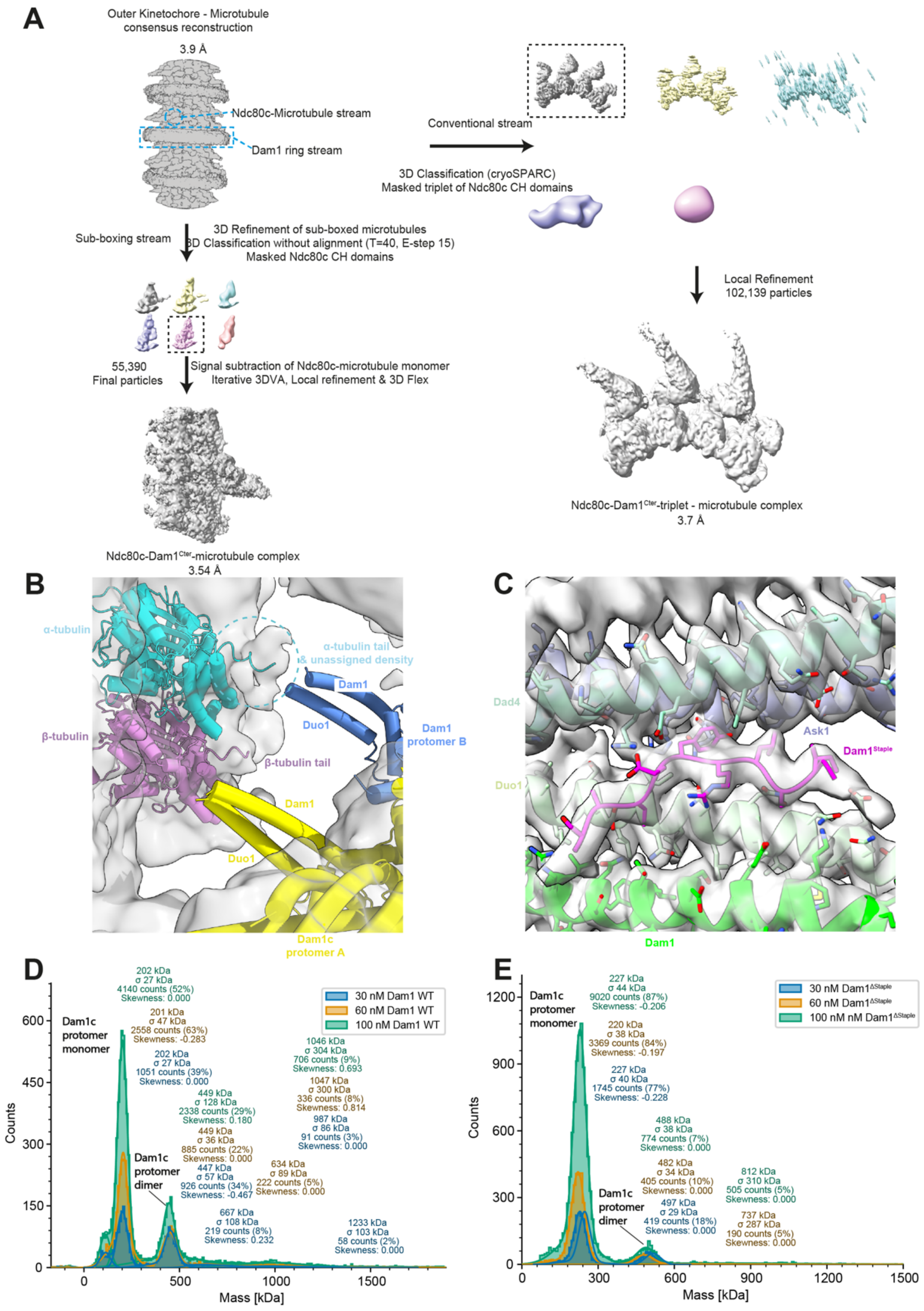


**Figure S2. Additional details of Dam1 structure determination, mass photometry experiments, and workflow for Ndc80c$^{CH}$:Dam1$^{C\text{-}ter}$ – microtubule focused cryo-EM processing. (A)** Zoomed view of map density beneath the Dam1 ring from the outer kinetochore microtubule consensus map shows diffuse density in the vicinity of the tubulin tails. **(B)** Map density for the Dam1$^{Protomer}$ dimer interface contoured around the corresponding molecular model to illustrate the maps can be interpreted at the level of amino-acid side-chains. Stacked histograms of mass photometry measurements taken of $Dam1^{Wt}$ **(C)** and $Dam1^{\Delta Staple}$ **(D)** at the indicated concentrations, with mass in kDa plotted against the number of observations. The molecular weights, error, and population of each mass class are indicated. At higher concentrations $Dam1^{Wt}$ forms higher order species that are essentially absent at corresponding titrations of $Dam1^{\Delta Staple}$ **(E)** Workflow for the Ndc80c:$Dam1^{C\text{-}ter}$ – microtubule cryo-EM processing stream.

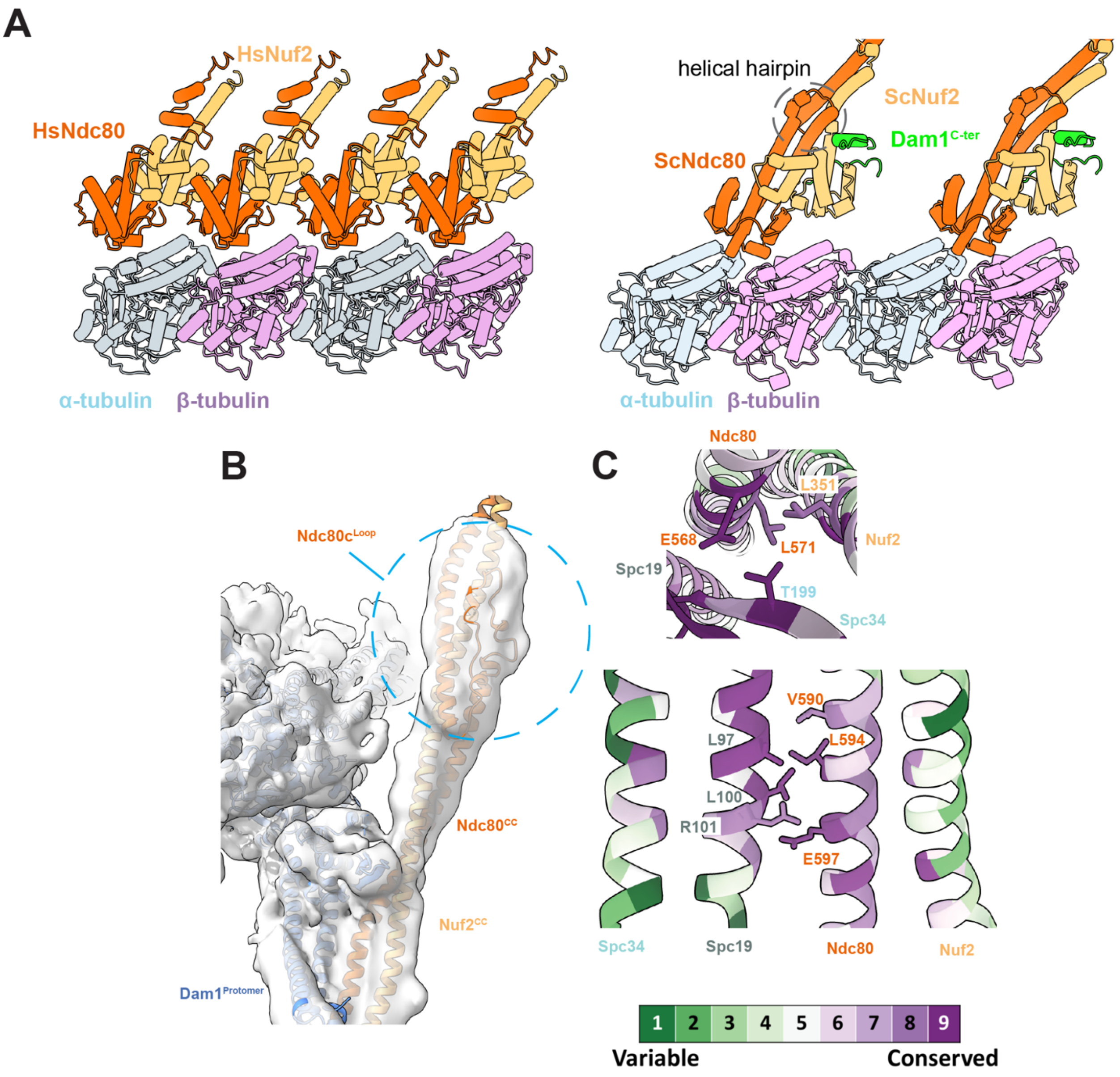


**Figure S3. Details of the Dam1c:Ndc80c and Ndc80c – microtubule interfaces. (A)** Side-by-side comparison of the mode of HsNdc80c$^{CH}$ (PDB: 3IZ0) and ScNdc80c$^{CH}$ binding to the microtubule. (**B**) Cryo-EM map corresponding to the Ndc80 loop region and with Ndc80 shown in cartoon representation. **(C)** Surface conservation representation of residues at the Ndc80-Nuf2 coiled-coil – Spc34-Spc19 interface. Interfacial residues are well conserved, whereas outward facing residues are not.

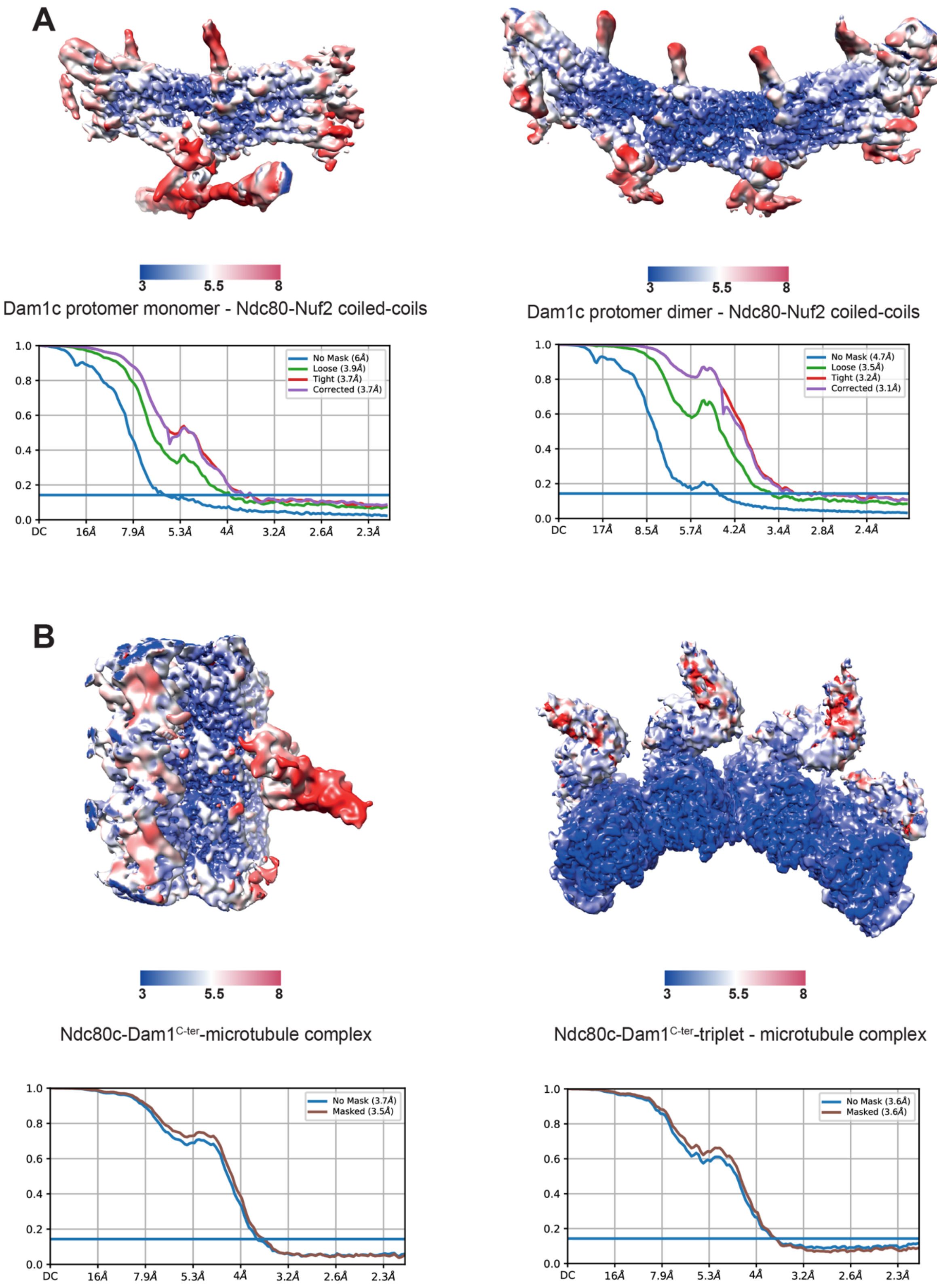


**Figure S4. Local resolution estimation and FSC curves for cryo-EM reconstructions. (A)** Local resolution estimation of Dam1 ring protomer – Ndc80-Nuf2 coiled-coil reconstructions (upper) and corresponding FSC curves (lower). The Dam1$^{Protomer}$ monomer – Ndc80$^{CC}$-Nuf2$^{CC}$ data are shown in the left panels, and the Dam1$^{Protomer}$ dimer – Ndc80$^{CC}$-Nuf2$^{CC}$ data on the right **(B)** Local resolution estimation of Ndc80c-Dam1$^{C\text{-}ter}$-microtubule reconstructions (upper) and corresponding FSC curves (lower). The Ndc80c$^{CH}$-Dam1$^{C\text{-}ter}$-microtubule data are shown in the left panels, and the Ndc80c$^{CH}$-Dam1$^{C\text{-}ter}$-microtubule triplet data on the right.

**Table S1. Cryo-EM data collection, refinement and validation statistics**

| | Ndc80c$^{CH}$- α/β-tubulin-Dam1$^{C-ter}$ (PDB XX) (EMDB XX) | Dam1c protomer dimer -Ndc80$^{CC}$-Nuf2$^{CC}$ (PDB XX) (EMDB XX) | Dam1c protomer monomer -Ndc80$^{CC}$-Nuf2$^{CC}$ (PDB XX) (EMDB XX) |
|---|---|---|---|
| **Data collection and Processing (for each dataset)** | | | |
| Microscope | TFS Titan Krios | TFS Titan Krios | TFS Titan Krios |
| Voltage (keV) | 300 | 300 | 300 |
| Camera | Gatan K3 | Gatan K3 | Gatan K3 |
| Magnification | 81,000 | 81,000 | 81,000 |
| Pixel size at detector (Å/pixel) | 1.06 | 1.06 | 1.06 |
| Total electron exposure ($e^{-}$/Å$^2$) | 50 | 50 | 50 |
| Exposure rate (e-/pixel/sec) | 16.6 | 16.6 | 16.6 |
| Number of frames collected during exposure | 50 | 50 | 50 |
| Defocus range (μm) | -1.2 to -3 | -1.2 to -3 | -1.2 to -3 |
| Automation software | EPU | EPU | EPU |
| Energy filter slit width (eV) | 20 | 20 | 20 |
| Micrographs collected (no.) | 39,610 | 39,610 | 39,610 |
| Micrographs used (no.) | 39,610 | 39,610 | 39,610 |
| Total extracted particles (no.) | 2,288,683 | 2,288,683 | 2,288,683 |
| **For each reconstruction:** | | | |
| Final particles (no.) | 95,730 | 248,732 | 77,272 |
| Point-group | C1 | C1 | C1 |
| Resolution (global, Å) | 3.54 | 3.15 | 3.97 |
| FSC 0.5 (unmasked/masked) | 4.9/4.5 | 8/4.2 | 7.8/7.4 |
| FSC 0.143 (unmasked/masked) | 3.7/3.5 | 4.7/3.1 | 4/3.9 |
| Resolution range (local, Å) | 2.29-30 | 2.29-30 | 2.275-30 |
| Map sharpening *B* factor (Å$^2$) | -104 | -60 | -85 |
| Map sharpening methods | CryoSPARC | CryoSPARC | CryoSPARC |
| **Model composition (for each model)** | | | |
| Protein (residues) | 1355 | 3298 | 1443 |
| Ligands | 4 | N/A | N/A |
| **Model Refinement (for each model)** | | | |
| Refinement package | PHENIX | PHENIX | PHENIX |
| - real or reciprocal space | Real Space | Real Space | Real Space |
| Model-Map scores | | | |
| -CC | 0.58 | 0.59 | 0.6 |
| *B* factors (Å$^2$) | | | |
| Protein residues (min/max/mean) | 45.83/115.47/92.61 | 30.00/1078.60/540.98 | 30.00/1078.60/524.10 |
| Ligands (min/max/mean) | 0.00/88.93/51.74 | N/A | N/A |
| R.m.s. deviations from ideal values | | | |
| Bond lengths (Å) | 0.004 | 0.003 | 0.008 |
| Bond angles (°) | 0.688 | 0.665 | 0.956 |

| | | | |
|---|---|---|---|
| **Validation (for each model)** | | | |
| MolProbity score | 1.88 | 1.96 | 2.29 |
| CaBLAM outliers | 0.98 (%) | 0.95 (%) | 1.01 (%) |
| Clashscore | 10.09 | 14.99 | 30 |
| Poor rotamers (%) | 0.17 | 0.26 | 0.67 |
| C-beta deviations | 0.00 | 0.06 | 0.65 |
| Ramachandran plot | | | |
| Favored (%) | 94.85 | 95.92 | 95.33 |
| Outliers (%) | 0.07 | 0.12 | 0.07 |

**Table S2.** Yeast strain genotypes

| |
|---|
| *MATa ade2-1 his3-11,15 trp1-1 leu2-3,112 can1-100 ura3-1::ADH1-OsTIR1(pMK200, URA3)* |
| *MATa ade2-1 his3-11,15 trp1-1 leu2-3,112 can1-100 ura3-1::ADH1-OsTIR1(pMK200, URA3),DAM1-mAID-FLAG$_3$::G418* |
| *MATa ade2-1 his3-11,15 trp1-1 leu2-3,112 can1-100 ura3-1::ADH1-OsTIR1(pMK200, URA3),DAM1-mAID-FLAG$_3$::G418, DAM1(Δ1-26)-V5$_3$::LEU2* |
| *MATa ade2-1 his3-11,15 trp1-1 leu2-3,112 can1-100 ura3-1::ADH1-OsTIR1(pMK200, URA3),DAM1-3xmAID-FLAG::G418, DAM1(Y17E, L19E, I21E)-V5$_3$::LEU2* |
| *MATa ade2-1 his3-11,15 trp1-1 leu2-3,112 can1-100 ura3-1::ADH1-OsTIR1(pMK200, URA3),DAM1-3xmAID-FLAG::G418, DAM1(I258A, L259A, I262A)-V5$_3$::LEU2* |
| *MATa ade2-1 his3-11,15 trp1-1 leu2-3,112 can1-100 ura3-1::ADH1-OsTIR1(pMK200, URA3),DAM1-3xmAID-FLAG::G418, DAM1(I258E, L259E, I262E)-V5$_3$::LEU2* |
| *MATa ade2-1 his3-11,15 trp1-1 leu2-3,112 can1-100 ura3-1::ADH1-OsTIR1(pMK200, URA3),DAM1-mAID-FLAG$_3$::G418, DAM1(Δ1-26, I258A, L259A, I262A)-V5$_3$::LEU2* |
| *MATa ade2-1 his3-11,15 trp1-1 leu2-3,112 can1-100 ura3-1::ADH1-OsTIR1(pMK200, URA3),DAM1-mAID-FLAG$_3$::G418, DAM1(Δ1-26, I258E, L259E, I262E)-V5$_3$::LEU2* |
| *MATa ade2-1 his3-11,15 trp1-1 leu2-3,112 can1-100 ura3-1::ADH1-OsTIR1(pMK200, URA3),DAM1-3xmAID-FLAG::G418, DAM1(Y17E, L19E, I21E, I258A, L259A, I262A)-V5$_3$::LEU2* |
| *MATa ade2-1 his3-11,15 trp1-1 leu2-3,112 can1-100 ura3-1::ADH1-OsTIR1(pMK200, URA3),DAM1-3xmAID-FLAG::G418, DAM1(Y17E, L19E, I21E, I258E, L259E, I262E)-V5$_3$::LEU2* |
| *MATa ade2-1 his3-11,15 trp1-1 leu2-3,112 can1-100 ura3-1::ADH1-OsTIR1(pMK200, URA3),NDC80-mAID-FLAG$_3$::G418* |
| *MATa ade2-1 his3-11,15 trp1-1 leu2-3,112 can1-100 ura3-1::ADH1-OsTIR1(pMK200, URA3),NDC80-mAID-FLAG$_3$::G418, NDC80 (E568A, V590A, L594A, E597A)-HA$_3$::TRP1* |
| *MATa ade2-1 his3-11,15 trp1-1 leu2-3,112 can1-100 ura3-1::ADH1-OsTIR1(pMK200, URA3),NDC80-mAID-FLAG$_3$::G418, NDC80 (E568R, V590W, L594E, E597R)-HA$_3$::TRP1* |

**Movie S1.**

Structure of the yeast outer kinetochore – microtubule complex, showing error correction phosphorylation sites at (i) the Dam1 staple, (ii) Ndc80c:Dam1$^{C\text{-}ter}$ interface (iii) Ndc80c:Spc34-Spc19, and finally a molecular model of the holokinetochore.
